# Mechanism of Nuclear Movements in a Multinucleated Cell

**DOI:** 10.1101/095695

**Authors:** Romain Gibeaux, Antonio Z. Politi, Peter Philippsen, François Nédélec

## Abstract

Multinucleated cells are important in many organisms but the mechanisms governing the movements of nuclei sharing a common cytoplasm are not understood. In the hyphae of the plant pathogenic fungus *Ashbya gossypii,* nuclei move back and forth, occasionally bypassing each other, and, preventing the formation of nuclear clusters, this is essential for genetic stability. These movements depend on cytoplasmic microtubules emanating from the nuclei, that are pulled by dynein motors anchored at the cortex. Using 3D stochastic simulations with parameters constrained by the literature, we predict the cortical anchors density from the characteristics of nuclear movements. Altogether, the model accounts for the complex nuclear movements seen *in vivo,* using a minimal set of experimentally determined ingredients. Interestingly, these ingredients power the oscillations of the anaphase spindle in budding yeast, but in *A. gossypii* this system is not restricted to a specific nuclear cycle stage, possibly as a result of adaptation to hyphal growth and multinuclearity.

## INTRODUCTION

Positioning the nucleus and mitotic spindle, appropriately within the cell, is critical in eukaryotes during many processes, ranging from simple growth to tissue development (Morris, 2002; Dupin and Etienne-Manneville, 2011; Gundersen and Worman, 2013; Kiyomitsu, 2015). Accordingly, cells have evolved various strategies to place their nucleus or spindle in the suitable location. In most studied systems, this process is driven by dynein pulling on nuclei-associated cMTs. Dynein is a minus-end directed motor that can act primarily in two ways. End-on pulling occurs when cortex-anchored dynein captures a growing cMT plus-end thereby initiating its shrinkage while maintaining the attachment. Lateral or side-on pulling occurs when cortex-anchored dynein walks on cMTs towards the minus end using its motor activity. This mode of action is independent of cMT shrinkage, and may result in the cMT sliding parallel to the cortex (Kotak and Gönczy, 2013; McNally, 2013; Akhmanova and van den Heuvel, 2016).

A detailed mechanistic view for directing nuclei during the cell cycle is known in the budding yeast *Saccharomyces cerevisiae* as outlined in Figure 1 and references therein. In contrast to most eukaryotic cells, the site of the cleavage plane in *S. cerevisiae* is selected early in the cell cycle by generating a ring structure (future bud neck) at the mother cell cortex at which the daughter cell (bud) will emerge. In addition, in budding yeast the nuclear envelope does not disassemble during mitosis, and nuclei are always attached to the minus-end of cMTs via spindle pole bodies (SPB) embedded in the nuclear envelope. The two pathways elucidated in *S. cerevisiae* involve cortical pulling but only the second pathway relies on dynein. During the Kar9-Bim1-Myo2 pathway, the nucleus is directed towards the bud neck by transporting cMT plus ends first along bud neck-emerging actin cables followed by depolymerization of the transported cMT. In metaphase, cMTs are transported along bud neck and bud tip-emerging actin cables securing positioning of the nucleus close to the bud neck and correct orientation of the spindle along the mother-bud axis. At pre-anaphase, when the bud has reached its final size, actin cables no longer emerge from the bud tip and the Dynein-Num1-pathway takes over. At the onset of anaphase, dynein, transported at the plus-end of cMTs to the bud cortex, starts pulling the spindle through the bud neck, as soon as dynein is captured by the cortical anchor Num1. An essential step in the cortical pulling is the switch from inactive dynein at cMT plus ends to active dynein after its association with Num1. This is regulated to happen first in the bud and later also in the mother cell explaining the observed back and forth movements of the anaphase spindle during mitosis in *S. cerevisiae.* The pulling of a cMT by “walking” of cortex-anchored dynein towards the cMT minus end at the SPB does not trigger depolymerization of this cMT, and the cMT slides head-on with its plus end along the cortex. Also, in contrast to higher eukaryotes, directing nuclei by dynein-mediated cortical pulling is restricted in *S. cerevisiae* to a small window of the cell cycle regulated by the inhibitor She1 (Woodruff *et al.*, 2009; Markus *et al.*, 2012).

**FIGURE 1:**
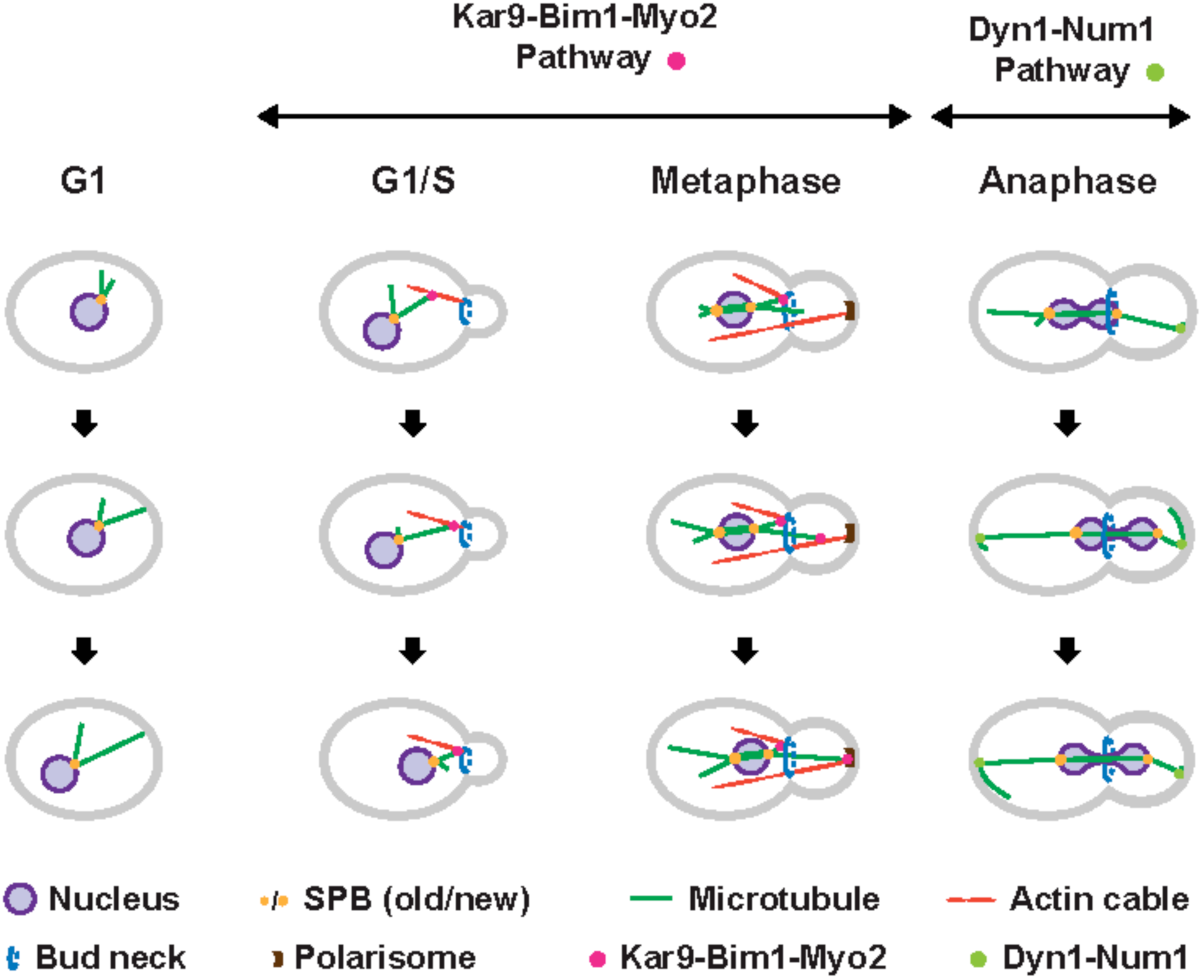
Nuclear movements in *S. cerevisiae.* Different modes of cMT +end capture and pulling on cMTs direct nuclei during the cell cycle of *S. cerevisiae.* **G1-phase:** cMTs arising from the SPB grow and shrink but more slowly than cMTs in animal cells (Carminati and Stearns, 1997). Nuclei can be pushed when growing cMTs hit the cortex (Shaw *et al.*, 1997). **G1/S-phase:** Actin cables emerge from the incipient bud site and the bud neck in small budded cells and are used as transport tracks by the type V myosin Myo2 (Pruyne and Bretscher, 2000). cMT +ends can be captured by Myo2, because Kar9 (APC tumor-suppressor homolog) connects the +end-tracking Bim 1 (EB1 homolog) and Myo2 indicated by the pink dot. Thus, the captured cMT is transported along actin cables to the bud neck, where shrinking of this cMT initiates pulling of the nucleus towards the bud neck, as necessary for nuclear positioning (Miller and Rose, 1998; Tirnauer *et al.*, 1999; Adames and Cooper, 2000; Yin *et al.*, 2000; Kusch *et al.*, 2002; Hwang *et al.*, 2003). **Metaphase:** After SPB duplication and separation, the old SPB stays close to the bud neck and accumulates Kar9 (Liakopoulos *et al.*, 2003). Cytoplasmic MTs emanating from this SPB can be captured, as described above, at their +end by Myo2 running at this cell cycle stage mainly along actin cables nucleated at the polarisome of medium- and larger-sized buds. Transport of captured cMTs along these cables secures the position of the nucleus and the correct orientation of the metaphase spindle. Anaphase: Prior to the onset of anaphase, dynein accumulates at the bud-proximal SPB, but dynein assumes a symmetric distribution at both SPBs soon after the pulling of the anaphase spindle through the bud neck is initiated (Grava *et al.*, 2006). Dynein is transported as an inactive complex with Pac1 (Lis1 homolog) at the +ends of cMTs (not shown), searching the bud cortex and later also the mother cell cortex (Markus and Lee, 2011; Markus *et al.*, 2011). When the immobile cortical Num1 captures dynein (light green dot), Pac1 is released and the activated dynein, still bound to the cMT, starts pulling the anaphase spindle through the bud neck (Lammers and Markus, 2015). The pulled cMT glides along the bud cortex. Frequently, the elongating anaphase spindle is pulled back into the mother cell (Yeh *et al.*, 1995; Adames and Cooper, 2000), most likely when dynein, captured and activated by Num1 at the mother cortex, exerts higher pulling forces on the mother cell SPB. This Dyn1-Num1 controlled oscillation of the anaphase pulling can repeat up to four times before the anaphase spindle is fully elongated.

The network of cMTs is also important in multinucleated cells (syncytia), which form at specific developmental stages in higher eukaryotes like the hundreds of nuclei in mammalian myotubes (Bruusgaard *et al.*, 2003) or the thousands of nuclei in fertilized eggs of insects (Foe and Alberts, 1983). During development of myotubes by cell fusions the cMT network is dramatically rearranged (Tassin *et al.*, 1985), the cMT plus end tracking protein EB3 regulates cMT dynamics at the cell cortex (Straube and Merdes, 2007), and nuclear movements depend on dynein and cMTs (Cadot *et al.*, 2012). Fertilized insect eggs initially develop in the absence of cytokinesis, and at later stages contain thousands of nuclei that eventually form a very organized layer near the cortex prior to the formation of cells around these nuclei. Recent progress in manipulating this system (Telley *et al.*, 2012) revealed that the spreading of nuclei depends on MT asters, F-actin and the elongating anaphase spindle. However, it is not exactly known how nuclei move apart and which cortical forces dictate their dispersion and final positioning at the cortex. An essential role of dynein and cMTs in controlling movements and distribution of multiple nuclei in a common cytoplasm has also been documented for several filamentous fungi (Morris, 2002; Xiang and Fischer, 2004; Fink *et al.*, 2006; Mouriño-Pérez *et al.*, 2006). Attempts to analyze molecular mechanisms for controlling movements of multiple nuclei in these fungi were hampered by the fact that dynein and cMTs are also involved in essential transports of organelles. A notable exception is the plant-pathogenic fungus *Ashbya gossypii,* which evolved from a budding yeast precursor based on its genome (Dietrich *et al.*, 2004, 2013) and which, like *S. cerevisiae,* exploits dynein and cMTs exclusively for nuclear movements in its multinucleated and constantly elongating cells, called hyphae (Figure 2A). Not only the genes but also cellular networks are highly conserved compared with *S. cerevisiae.* Thus, the different cellular life style of *A. gossypii* is driven by homologs of well-studied *S. cerevisiae* genes, in particular those implicated in the cell cycle, polar growth and cytoskeleton. With respect to *A. gossypii* nuclear biology, we begin to fully understand asynchronous mitoses of multiple nuclei in a common cytoplasm (Gladfelter *et al.*, 2006; Lee *et al.*, 2013), but we are still missing a comprehensive mechanism for the complex nuclear motility during hyphal growth.

**FIGURE 2:**
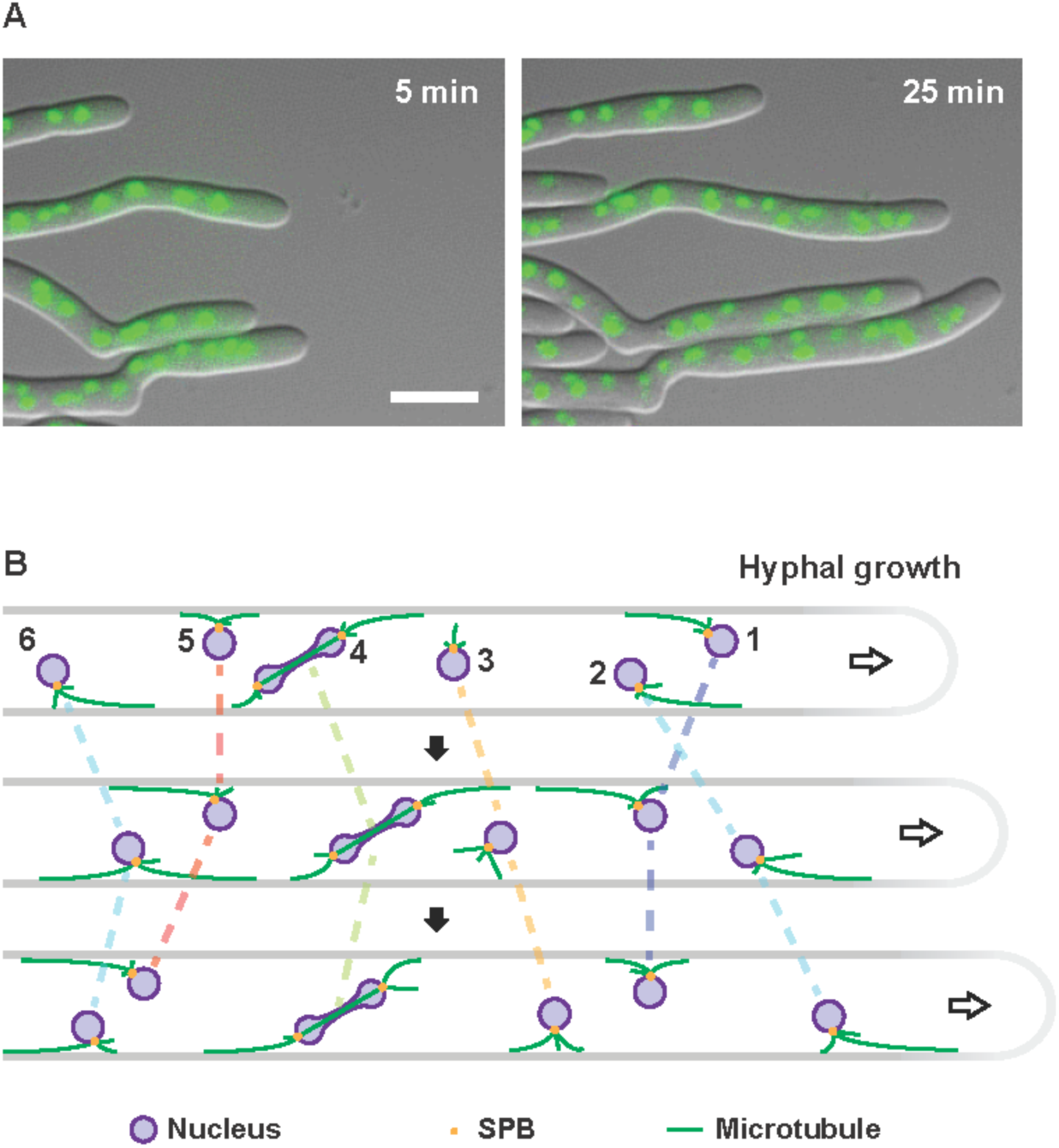
Nuclear movements in *Ashbya gossypii.* (A) Visualization of long range nuclear migration and nuclear dynamics in hyphae of *A. gossypii.* Two merged DIC and fluorescence images of hyphae with histone H4-GFP marked nuclei of Video 1 were selected to show the efficient polar growth and maintenance of nuclear density. The bar represents 10 μιη. (B) Qualitative model of nuclear migration control in a hypha growing from left to right which causes a cytoplasmic flow (arrow) in the growth direction. The indicated nuclei do not necessarily represent the most apical six nuclei. Nuclei are autonomous with respect to movements and mitosis (Alberti-Segui *et al.*, 2001; Gladfelter *et al.*, 2006; Lang *et al.*, 2010b). Each spindle pole body (yellow dots) nucleates 3 cMTs (green lines) based on electron tomography (Gibeaux *et al,* 2012). Very long cMTs can form because the dynamic cMT parameters are adapted to the very long cells (Grava and Philippsen, 2010; Lang *et al,* 2010b). Like in *S. cerevisiae,* dynein localizes at the growing end of cMTs (Grava *et al.*, 2011), and Num1 patches form at the cortex (Schmitz and Landmann, unpublished). Therefore, dynein can be captured at the cortex by Num1 which initiates pulling on the cMT thereby moving the nucleus, essentially as described for pulling of the anaphase spindle in *S. cerevisiae* by the dynein off-loading model (Lee *et al.*, 2003, 2005; Sheeman *et al.*, 2003; Markus and Lee, 2011; Lammers and Markus, 2015). The cMT continues growing during the pulling (Grava and Philippsen, 2010) until it eventually will start depolymerizing if its +end undergoes a catastrophe. **Nucleus 1** is pulled backward and **nucleus 2** forward. The distances pulled are sufficiently long to cause nuclear bypassing. Next, nucleus 1 stops moving due to balance of backward and forward pulling forces while nucleus 2 continues to move forward. **Nucleus 3** migrates with the cytoplasmic flow. No cortical contact of cMTs. **Nucleus 4** first moves forward and then backward as the mitotic nucleus is pulled. Up to a certain length, elongating spindles can assume any angle with respect to the growth axis (Alberti-Segui *et al.*, 2001; Grava *et al.*, 2011). **Nucleus 5** is immobile first, then moves backward because the backward-oriented cMT elongated and is pulled. **Nucleus 6** moves forward and switches to backward movement because the forward-oriented cMT shrinks and the backward-oriented cMT grows, changing the balance of forces.

Within the hyphae of *A. gossypii,* nuclei permanently perform short-range back and forth excursions, which presumably help avoiding formation of nuclear aggregates, as well as longer-range movements, frequently leading to nuclear bypassing events that permute the order of the nuclei (Alberti-Segui *et al.*, 2001; Gladfelter *et al.*, 2006; Grava *et al.*, 2011). This continuous reordering prevents that a detrimental mutation acquired in the haploid genome of one nucleus would remain concentrated in a hyphal segment, compromising it. Nuclear movements require cMTs and functional SPBs permanently embedded, like in *S. cerevisiae,* in the nuclear envelope. In the absence of cMTs or functional SPBs, oscillatory and bypassing movements do not occur and nuclei are passively transported with the cytoplasmic stream in the direction of hyphal tip growth (Alberti-Segui *et al.*, 2001; Gladfelter *et al.*, 2006; Lang *et al.*, 2010a, 2010b). *A. gossypii* cMTs grow with 6 μm/min, three times faster than *S. cerevisiae* cMTs, and thus can always reach hyphal tips the maximal speed of which is 3.3 μm/min (Grava and Philippsen, 2010).

Interestingly, a single type of minus-end directed MT motor – dynein – is responsible for all active nuclear movements during hyphal growth. Indeed, reducing the expression level of the dynein heavy chain by truncating the AgDYN1 promoter led to the decrease of oscillatory and bypassing movement frequencies (Grava *et al.*, 2011). Dynein is localized at different cMT positions: At cMT minus ends (SPBs), along cMTs and at cMT plus ends, especially of long cMTs. It has been suggested that, once anchored to the hyphal cortex, dynein could exert pulling forces that would move the nuclei (Grava and Philippsen, 2010), essentially as proposed for the anaphase spindle pulling in *S. cerevisiae* (Lee *et al.*, 2003; Sheeman *et al.*, 2003). This is supported by the phenotype of hyphae lacking functional AgNum1 (Grava *et al.*, 2011). In the same study, it was shown that AgKar9 or AgBim1 are not important for bidirectional movements and bypassing of nuclei, which is not surprising because nuclear positioning is not an issue in growing *A. gossypii* hyphae. Electron tomography revealed that each SPB nucleates on average 3 cMTs and that SPBs are frequently found as duplicated side-by-side entities (G2 phase of the nuclear cycle) (Gibeaux *et al.*, 2012). This is in line with the *in vivo* observation that up to 6 cMTs emanate from one nucleus (Lang *et al.*, 2010b).

Altogether, these observations suggest a mechanistic hypothesis for nuclear motility in *A. gossypii* hyphae: Highly dynamic cMTs emanating in different directions from the nuclear envelope-embedded SPB would reach the cortex, and be pulled by dynein anchored to it (Figure 2B). This model is appealing because it involves only one motor and seems qualitatively complete. Yet, it is not clear if a single mode of action could explain the complexity of the observed movements. A detailed quantitative analysis of the mechanism using simulations is thus essential to reveal if certain features of nuclear positioning are not correctly captured by the current proposed model, and if this model rather needs to be refined or investigated further through additional experiments. To this end, we implemented this model with *Cytosim* (www.cytosim.org), to simulate the motion of multiple nuclei in a 3D geometry, using flexible and dynamic cMTs. Using physically realistic equations of motion, the simulation computes the nuclei movements that can be directly compared to the movements observed in live-imaging experiments. With parameter values obtained from published experimental measurements, our simulation was able to match the basic characteristics of nuclei movements, and we could thus test different experimental settings relating to how cMT dynamics, cMT number per nucleus, cortical Num1 density, cytoplasmic streaming or organelle crowding impact on nuclear movements in multinucleated hyphae.

## RESULTS

### A computational method for validation of natural and simulated nuclear movements

The nuclei in live *A. gossypii* hyphae can easily be observed with fluorescence microscopy (Figure 2A; Video 1), and their bidirectional movements including bypassing have already been characterized (Alberti-Segui *et al.*, 2001; Gladfelter *et al.*, 2006; Grava and Philippsen, 2010; Lang *et al.*, 2010b; Grava *et al.*, 2011; Anderson *et al.*, 2013). In these publications, different methods were employed to measure nuclear movements, most not applicable for our computational simulation. Therefore, we established an automatic method to measure nuclear motility parameters which we first applied to re-analyze *in vivo* nuclear movements. Spatial and temporal information was extracted from videos of 12 hyphae expressing GFP-tagged histone H4 recorded every 30 s over 25-30 min (see example in Figure 2B and Video 1). In each hypha, we tracked the 2D positions of the first 5 nuclei (and in few cases their progeny) to accurately follow their movements. The dataset was rotated to calculate the position of the nuclei along the hypha main axis. A global analysis, first, showed that nuclei are moving in the direction in which the hyphae grows (Supplemental Figure S1A), at a speed that matches the growing speed for nuclei located near the tip of the hyphae. We thus used the measured growth speed of the hyphal tip to set the flow in the simulation. On top of this streaming motion, the analysis showed that cMT-dependent nuclear motility is diffusive at long time scales, with an effective diffusion coefficient of ~0.0113 μm^2^/s (Supplemental Figure S1B). By comparison, the Brownian motion of the nuclei leads to a diffusion coefficient of ~0.0003 μm^2^/s, using Stokes’ law with our estimate of cytoplasmic viscosity. Hence, although cMT-driven motions of nuclei appear non-directed, they increase the diffusion speed dramatically. We also expect that Brownian forces alone would not be sufficient to permute the order of the nuclei such that nuclear bypassing is a hallmark of cMTs dependent movements.

We then performed a more detailed analysis, as illustrated in the first three panels of Figure 3 for one representative hypha. We first generated a standard representation of patterns of nuclear movement to allow visual comparisons to previously published analyses (Figure 3A) and then produced standardized schemes to document the frequent changes in the direction and speed of individual nuclei (Figure 3B and C, respectively). Using the cytoplasmic flow speed *v_h_* as a threshold, we classified the movements in three categories: forward (*v_h_* < *v*), backward (*v* < 0) and tumbling (0 < *v* < *v_h_*). From this, we computed frequencies, durations and average speed (i.e. average of the instantaneous velocities) for each class. The result of the analysis of the 12 hyphae (70 nuclei) is reported in Supplemental Table S1, and the average of each motility parameter in Table 1. As can be seen in the top row of this table, on average nuclei performed 0.279 (± 0.033 SD) forward movements per minute and 0.208 (± 0.065 SD) backward movements per minute with an average speed of 0.864 (± 0.209 SD) μm/min. The average ratio of forward to backward frequencies (here 1.628 ± 1.190 SD) will be used later extensively for comparing simulation conditions, because it is a simple yet sensitive readout of the movements. We also automatically detected nuclei reordering events – when one nucleus bypasses another one. These events occur at a frequency of 0.026 (± 0.023 SD) per minute which means that about 10% of the forward or 8% of backward movements lead to nuclear bypassing.

**FIGURE 3:**
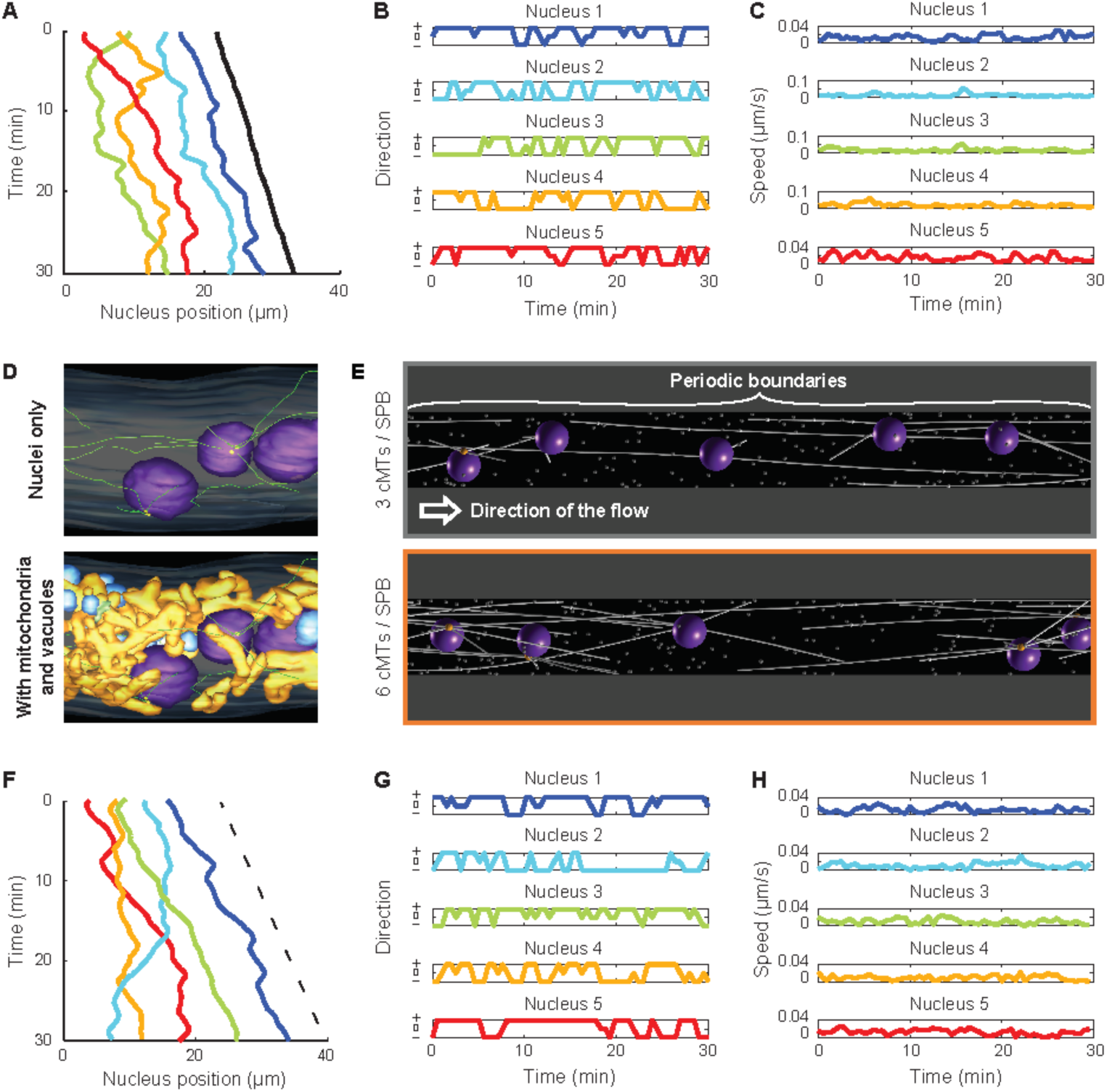
Analyzing in vivo and simulated nuclear movements. **(A)** The position of five nuclei along time showing movement patterns. The center of individual nuclei (1: Blue, 2: Turquoise, 3: Green, 4: Orange, 5: Red) and the position of the hypha tip (Black) were tracked along the course of the movie. Three reordering (bypassing) events can be seen, whereby two nuclei exchange their ranking along the hyphae. **(B)** The category of movement of each nucleus as a function of time. From the coordinates of each nucleus (1-5), the signed axial speed of movement *v_x_* was computed at every time point. The hypha growth speed *v_h_* was used as a threshold to categorize the movements as follows: forward ‘+’ (*v_h_* < *v_x_*), backward ‘−’(*v_x_* < 0) and tumbling ‘0’ (0 < *v_x_* < *v_h_*). Colors match panel A. **(C)** Plots of absolute nuclei speed along time. Magnitude of nuclei speed along time (||*v*||) does not distinguish between forward and backward directions. Colors match panel A. **(D)** Snapshots of electron tomography models. Previously publish data (Gibeaux *et al.*, 2012, 2013) were used to generate snapshots showing on the top panel cMTs (green) nucleation and organization from nuclei (purple) - embedded SPBs (yellow) and on the bottom panel, the same nuclei within the cytoplasm crowd, including mitochondria (orange) and vacuoles (blue). **(E)** Snapshot of simulations. The simulations are performed in 3D within a cylinder with periodic boundaries in x, as indicated. The simulations contain 5 nuclei (dark violet) and each nucleus nucleates 3 (top) or 6 (bottom) cMTs (white lines). Cytoplasmic MTs are attached at their minus ends to one point at the surface of the nucleus (the SPB, orange ball) and grow at their plus ends. Inactive minus end-directed dynein motors are assumed to be present at each plus end. Each contact of a cMT plus end with a cortical dynein anchor (white dots) will induce pulling. The arrow indicates the direction of the applied cytoplasmic flow. **(F)** Nuclear positions along time taken from a simulation with 6 cMTs per nucleus. The *x*-coordinates of the center of individual nuclei are plotted in various colors (1: Blue, 2: Turquoise, 3: Green, 4: Orange, 5: Red), while the dashed line indicates the applied flow speed. **(G)** The direction of motion of each nucleus along time. The motions were categorized exactly as in panel B. Colors match panel F. **(H)** Plots of absolute nuclei speed along time, exactly as in panel C. Colors match panel F.

**Table 1.**
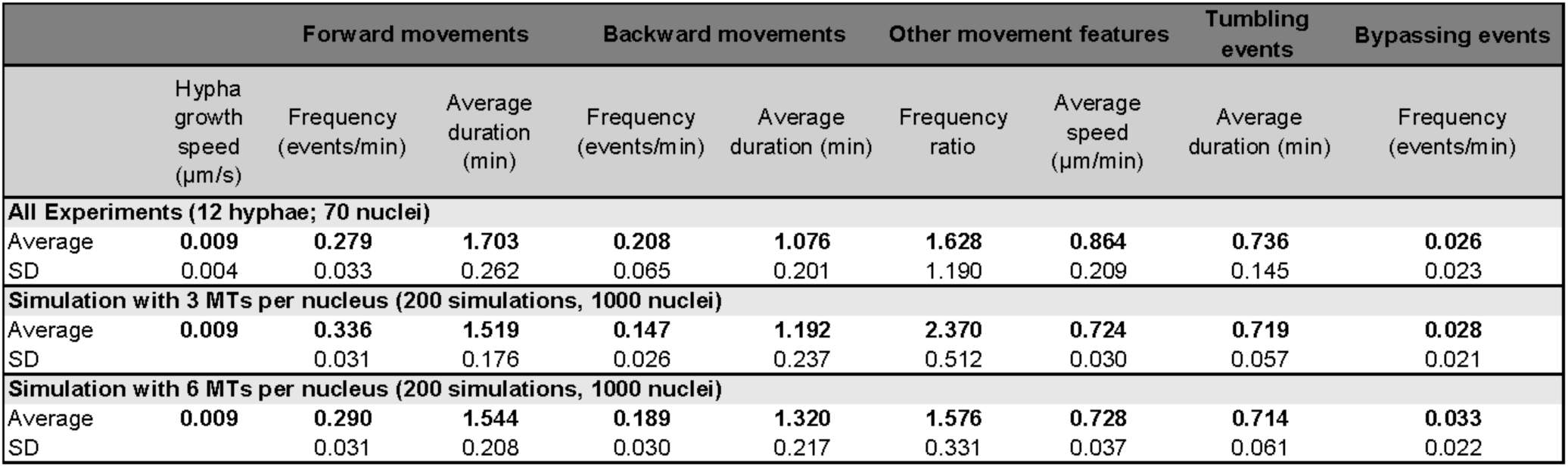
Nuclear dynamics quantification averages from live-imaging experimental data and *in silico* experiments.

### Implementation of a simulation system for nuclear movements in *A. gossypii*

Accumulated evidence summarized in the introduction and the legend of Figure 2B supports a qualitative model for nuclear movements in *A. gossypii:* Cortex-anchored dynein exert pulling forces on dynamic cMTs, nucleated from SPBs, and thus drive the movements of nuclei at all cell cycle stages (Figure 2B). To test whether this model is physically plausible and sufficient to explain the observed complex nuclear motility, we generated a computer simulation of the process within hypha-like 3D geometry using the *Cytosim* platform (Nedelec and Foethke, 2007). For cell biological parameters of the simulation, we considered numerous life cell imaging data mentioned above. We also considered high-resolution EM studies including the EM tomography-based 3D model of an *A.gossypii* hypha (Lang *et al.*, 2010b; Gibeaux *et al.*, 2012, 2013). For example, nuclei with one SPB emanate 3 cMTs but a high percentage of nuclei carry two side-by side SPBs emanating 6 cMTs (Figure 3D, top image). We therefore implemented simulations with 3 cMTs and simulations with 6 cMTs per nucleus for each tested condition. To account for the organelle-crowded cytoplasm, we simulated mitochondria and vacuoles (Figure 3D, bottom image, orange and blue organelles, respectively) as spheres the number and volume of which matched the measurements as discussed in more detail below Figure 7). In all simulation images, these spheres are present but not displayed for clarity.

We knew that 30 min monitoring of 5 nuclei in a 30 μm long hyphal segment, corresponding to the average *in vivo* spacing of nuclei (Lang et al., 2010a), is sufficient to document nuclear motility parameters, and we therefore simulated a hyphal segment of this size as a cylindrical volume with periodic boundaries. In the graphical representations of simulated hyphae shown in Figure 3E, a nucleus or a cMT may thus leave the cylinder at the right side and reappears at the left side, or *vice versa.* There is, however, no edge in the simulated space. The two simulation images of Figure 3E display the elements that were implemented to test whether a nuclear motility pattern similar to wild type hypha can be reproduced. All cMTs (thin white rods) emanate from one site (SPB or duplicated SPB) of a nucleus, they remain attached to this site while they are elongating, they can undergo very fast shortening until they disappear inducing at this SPB a new cMT to form immediately. Cytoplasmic MTs exclude each other and cannot penetrate nuclei and other simulated organelles, they can elongate in all directions and, upon contact of a tip (+end) with the cylinder boundary (cortex), continue to elongate while sliding along the cylinder boundary. They are programmed to engage in pulling forces (thereby moving the attached nucleus) when they hit one or more of the randomly distributed immobile white dots at the simulated cortex (patches of the dynein-activating anchor Num1) highlighted by increased dot intensity, and they are programmed to terminate pulling according to a detachment rate and in response to counteracting forces. All quantitative parameters used for implementing the simulation of nuclear movements, such as the viscosity of the cytoplasm, were obtained from the literature and are reported in Table 2, with the exception of the fitted cortical density of Num1 (see Figure 5).

**FIGURE 4:**
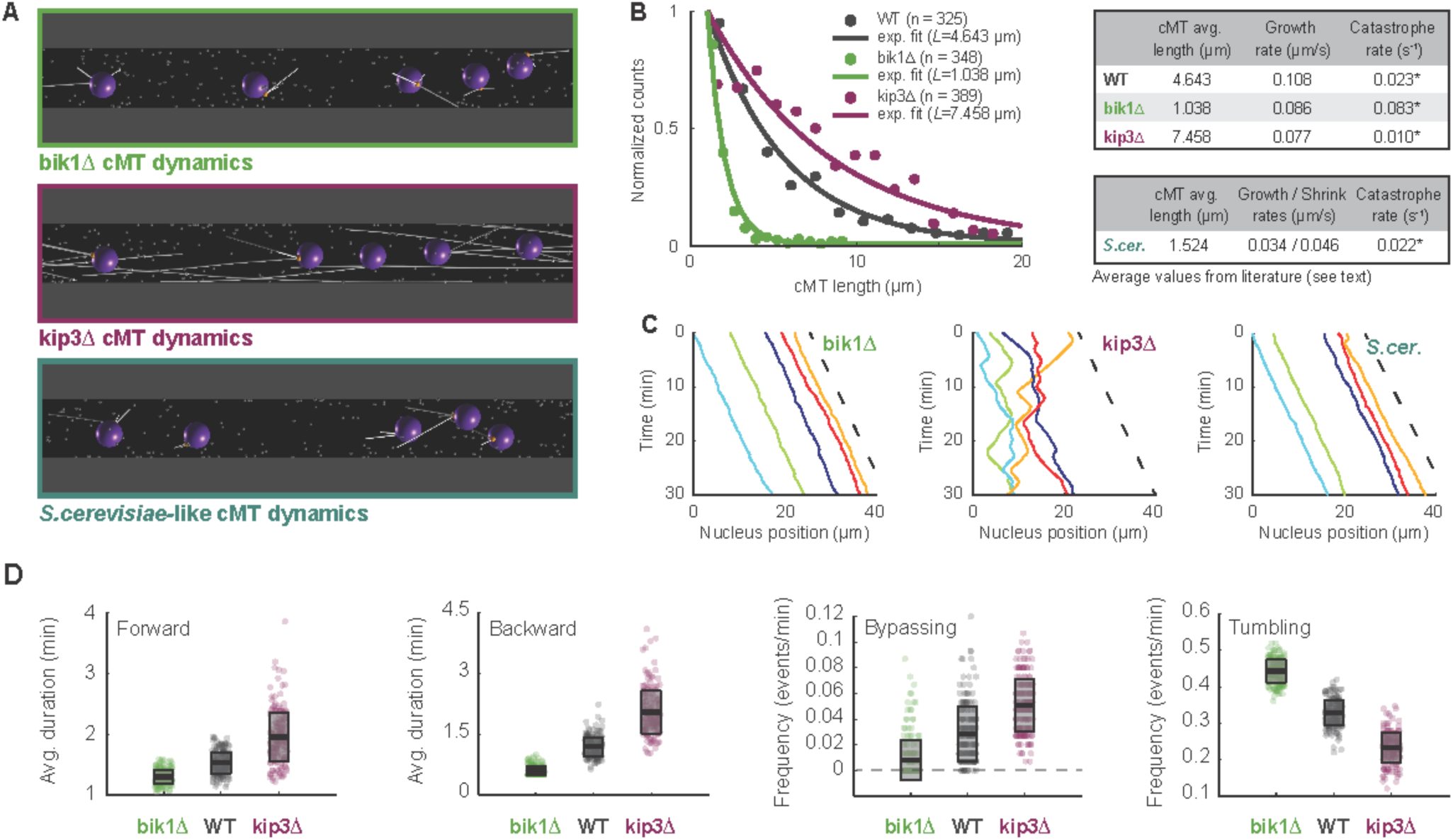
Verification of nuclear mobility phenotypes in mutants with altered cMT dynamics. **(A)** Snapshot of simulations with mutant cMT dynamics. The bikl Δ (top; green), kip3 Δ (middle; magenta) and *S. cerevisiae-like* (bottom; dark turquoise) conditions are shown. **(B)** Distributions of cMT length in WT, bik l Δ and kip3Δ cells. Because, as shown by electron tomography, many cMTs are below the light microscopy resolution, only cMT length values (Grava and Philippsen, 2010) from 1 to 20 μιη were used (*n* corresponds to the number of data point in this range). The frequency of each bin (circles) and an exponential fit on the truncated distribution (line) is shown (Fraile and García-Ortega, 2005). The corrected average cMT length *L* was determined. Data for WT are plotted in gray, for bik l Δ in green and for kip3Δ in magenta. Tables summarize the determined catastrophe rates for WT, bik l Δ, kip3Δ and the *S. cerevisiae-like* mutants. The catastrophe rates were calculated (*) by dividing the published cMT polymerization rates *v_g_* by *L* (see Results and Materials and Methods). **(C)** Plots of nuclear position along time in b i kl Δ (left), ki p3 Δ (middle) and *S. cerevisiae-like* (right) simulations. The dashed line indicates the applied flow speed. **(D)** Boxplots of nuclear dynamics measurements in “wild-type” and “mutant” simulations. The average duration of forward movements (left), the average duration of backward movements (middle left), the frequency of bypassing events (middle right) and the frequency of tumbling events (right) are plotted. Simulations representing ‘biklA’, ‘WT’ and ‘1<ρ3Δ’ cells are plotted in green, gray and magenta, respectively. Circles stand for individual simulations; the thick black line marks the average value and the transparent gray box the standard deviation.

**FIGURE 5:**
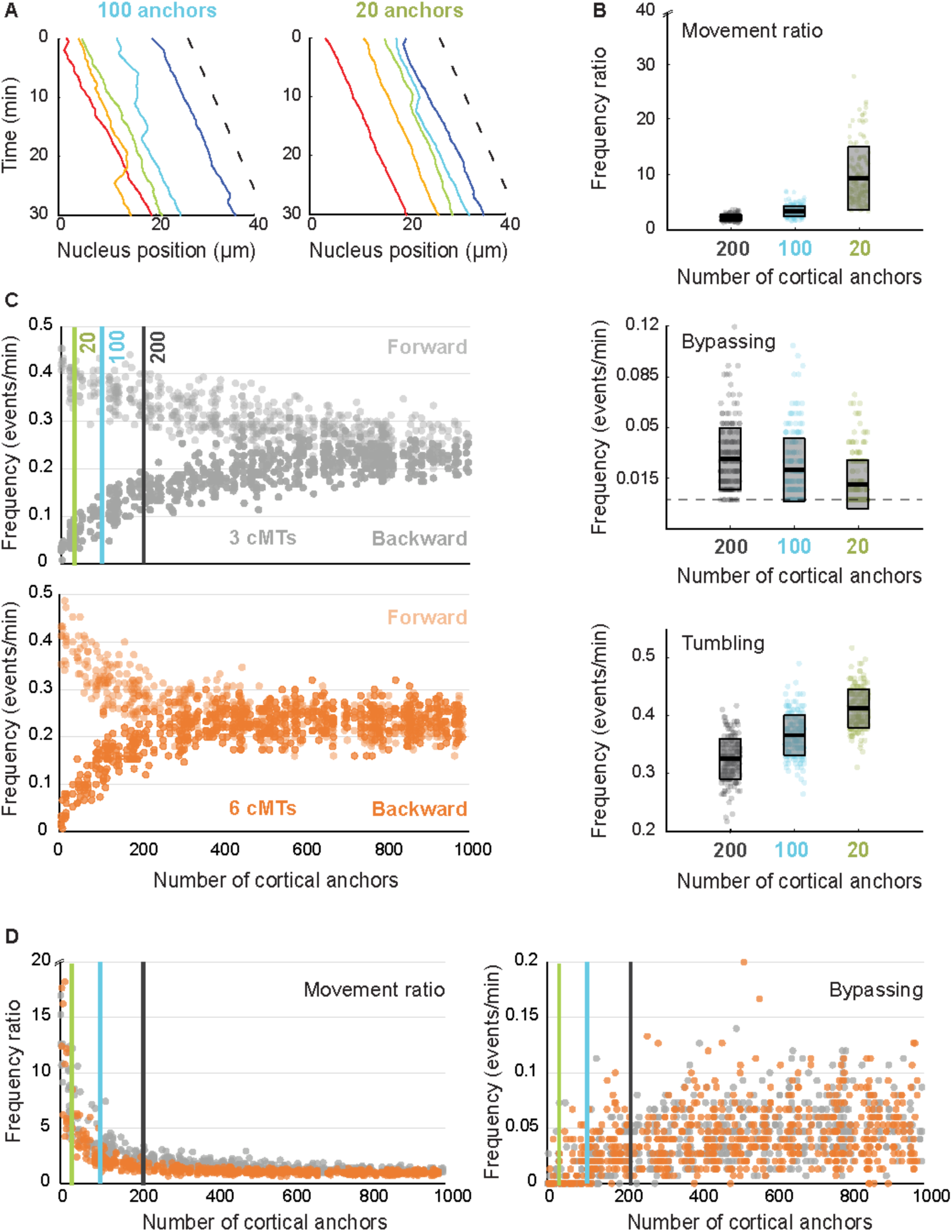
Importance of anchor density at the cell cortex. (A) The position of five nuclei along time in simulations containing 100 (left) and 20 (right) cortical anchors per 30 μm of hypha. The dashed line indicates the applied flow speed. (B) Quantification of nuclear movements in simulations with 200 (“wild-type”), 100 and 20 cortical anchors. The ratio of forward / backward movement frequencies (top), the frequency of bypassing events (middle) and the frequency of tumbling events (bottom) are shown. The dots stand for individual simulations, and the box indicates the mean and standard deviations. Results obtained with 200, 100 and 20 anchors are plotted in gray, turquoise and lime green, respectively. (C) The frequencies of forward (lighter) and backward (darker) movements as a function of the number of anchors, with 3 cMTs (gray; top) or 6 cMTs (orange; bottom) per nucleus. Each vertical line at 20, 100 and 200 marks the position of the parameter values used in B. (D) The ratio of forward / backward movement frequencies (left) and the frequencies of bypassing events (right) as a function of anchor density for 3 cMTs (gray) and 6 cMTs (orange). The three vertical lines mark the position of the parameter values used in B.

**Table 2.**
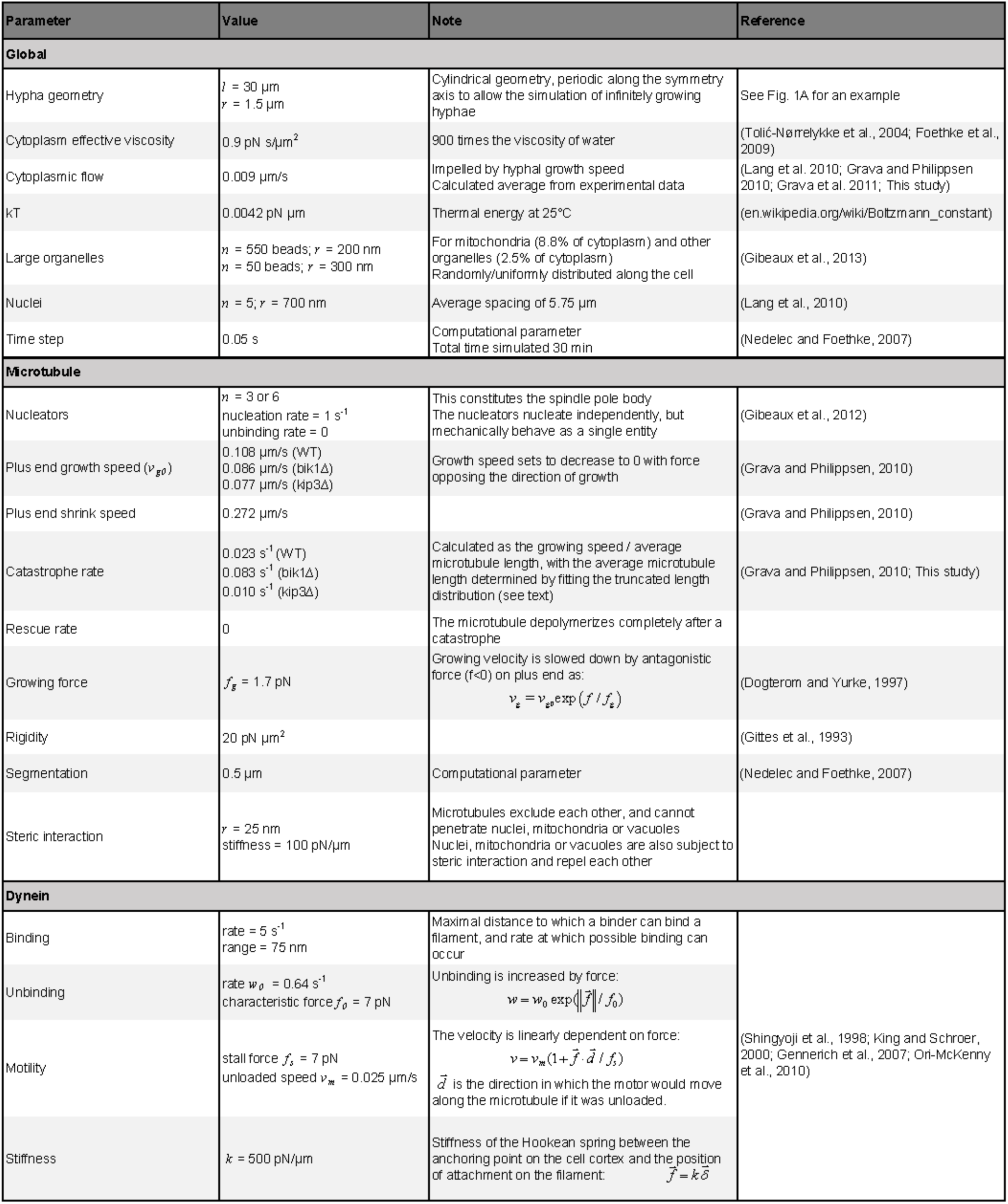
Parameters of the simulation

We also created a flow component to model the effect of the cytoplasmic streaming present in growing hyphae. Unlike in fungal colonies such as the ones made by *N. crassa* (Ramos-García *et al.*, 2009; Roper *et al.*, 2013), the hyphae of *A. gossypii* do not fuse with each other, and the flow does not change directions. The flow is thus characterized by a speed *𝑣_flow_* (Figure 3E), and for simplicity, we neglected the detailed micro flows that could arise around nuclei, and their hydrodynamic interactions, and only adjusted the viscous drag force in the equation of motion to: — *ξ*( *𝑣 — 𝑣_flow_*), where *ξ* is the viscous drag coefficient. The flow affects all objects in the simulation, such that any object moves at the speed of the flow, unless it is subjected to other forces than the drag. The average cytoplasmic streaming in our *in vivo* dataset was 0.009 μm/s (Table 1), and we used this value for the parameter *𝑣_flow_* in most simulations. The effects of much lower or faster cytoplasmic flow speeds on nuclear movements are discussed below (see Figure 6).

**FIGURE 6:**
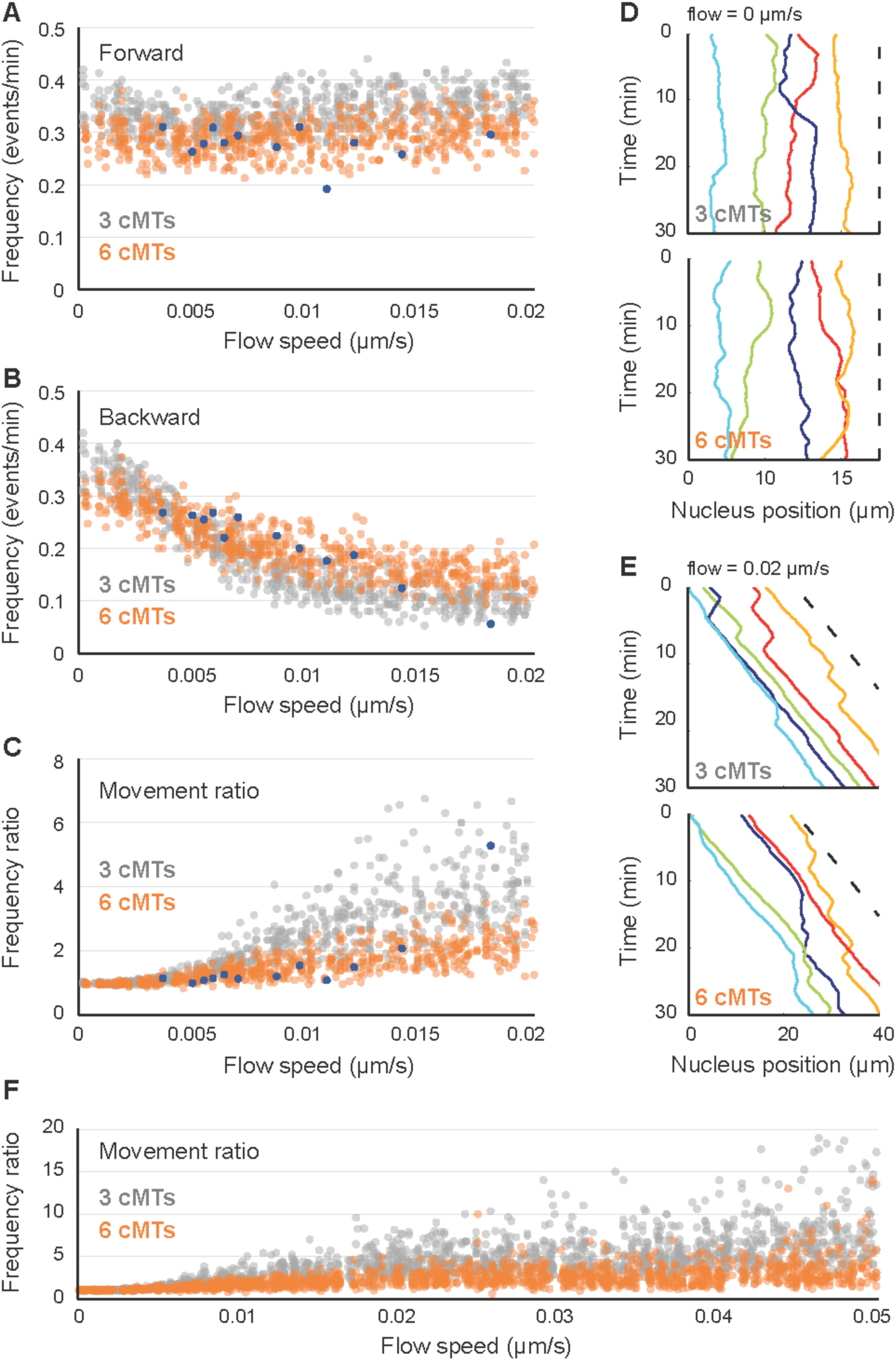
Influence of cytoplasmic flow on nuclear movements and role of cMT number in movement balance. **(A)** The frequency of forward movements as a function of increasing cytoplasmic flow (from 0 to 0.02 μm/s). Simulation values are plotted in gray (3 cMTs per nucleus) and orange (6 cMTs per nucleus) and experimental values in blue. **(B)** Plot of the frequency of backward movements as a function of increasing cytoplasmic flow (from 0 to 0.02 μm/s). Colors as in panel A. **(C)** The ratio of movement frequencies (forward/backward) as a function of increasing cytoplasmic flow (from 0 to 0.02 μm/s). Colors as in panel A. **(D)** Simulated nuclear positions along time without flow (speed of 0 μm/s). The *x*-coordinates of the center of individual nuclei (1-5) is plotted in different colors, while the dashed line indicates the applied flow speed. Top is an example of a simulation with 3 cMTs. Bottom is an example of a simulation with 6 cMTs. **(E)** Simulated nuclear positions along time with a flow speed of 0.02 μm/s. Same legend as D. **(F)** The ratio of movement frequencies (forward/backward) as a function of increasing cytoplasmic flow (from 0 to 0.05 μm/s). Colors as in panel A.

### *A. gossypii* nuclear movement patterns can be reproduced *in silico*

We computed 200 simulations (1000 nuclei) with 3 cMTs nucleated per nucleus (Video 2), and another set with 6 cMTs per nucleus (Video 3). To provide a visual readout and for comparison purpose, we plotted the patterns of nuclear movement for one simulation with 6 cMTs per nucleus (Figure 3F to H). The short-range oscillations and long-range movements as well as the patterns of direction changes and the speed of movements were very similar as the *in vivo* measurements (compare to Figure 3A to C). This similarity also holds when the averages of all simulations are compared with the *in vivo* measured values (Table 1). With 3 cMTs per nucleus, the frequencies of movements were: forward 0.336 (± 0.031 SD) min^−1^, backward 0.147 (± 0.026 SD) min^−1^, giving a ratio of 2.370 (± 0.512 SD) (Table 1). With 6 cMTs per nucleus, backward movements were more pronounced: forward = 0.290 (± 0.031 SD) min^−1^, backward = 0.189 (± 0.03 SD) min^−1^, with a ratio of 1.576 (± 0.331 SD) (Table 1). Interestingly, the simulations with 6 cMTs per nucleus matched the values extracted from experimental data better, which is consistent with the previous observation that *A. gossypii* hyphae contain more nuclei with duplicated side-by-side SPBs than with a single SPB (Gibeaux *et al.*, 2012). Surprisingly, the change in cMT number does not seem to influence the frequency of bypassing events, although it affects the balance between forward and backward movements (Table 1).

Finally, it was previously observed that the SPB is leading the nuclear mass during a pulling event. If the direction of pulling changes, the SPB relocates within a minute to the opposite side of the nucleus, which becomes the new leading front (Lang et al., 2010a). These detailed SPB movements were actually well captured in the simulation (see Video 4).

### Phenotypes of mutations affecting microtubule dynamics are verified by simulations

Nuclei in hyphae lacking Bik1, the cMT +end stabilizing homolog of CLIP170, generate shorter cMTs compared to wild type, and show decreased frequencies of bidirectional movements and no bypassing. The growth of this deletion mutant is not affected because nuclei migrate with the cytoplasmic stream. In contrast, nuclei in hyphae lacking Kip3, the cMT destabilizing homolog of kinesin-8, generate longer cMTs and display increased frequencies of bidirectional movements and bypassing compared to wild type (Grava and Philippsen, 2010). We wanted to know to which degree simulations with altered cMT dynamic are able to reproduce known changes in nuclear mobility patterns caused by mutations affecting cMT dynamics. We thus decided to implement, in a new set of simulations, parameters representing the changed cMT dynamics in the bik1 and kip3 mutant. In addition, we implemented in a third set of simulations the dynamic instability parameters of *S. cerevisiae* cMTs.

Graphical images representing the cMT length differences in simulations with altered cMT dynamic are shown in Figure 4A. The parameters for dynamic instability of cMTs in hyphae lacking Bik1 or Kip3 were modeled using the “classical” two-states model (Figure 4B). For this modeling, growth and shrinkage speeds are constant, and catastrophes occur stochastically as first-order events. Rescues are not considered, because rescues have not been observed in *A. gossypii,* but new cMTs are nucleated by the SPB at each unoccupied site. We used the cMT polymerization rates *vg* measured in each mutant (Grava and Philippsen, 2010) and estimated the catastrophe rate from the lengths of cMTs that had been determined by light microscopy of fixed and immuno-labeled mutant hyphae. Specifically, we estimated the average length *L* by fitting an exponential on the truncated length distributions, between 1 and 20 μm (Figure 4B and Materials and Methods). Unlike the average of all values, this procedure (Fraile and García-Ortega, 2005) avoided the bias in the data due to the fact that shorter cMTs would be missed in light microscopy. The longest cMT in the WT dataset was 26.6 μm. and 20 μm was chosen as an upper limit to include most of the data in all conditions. The catastrophe rate was then calculated as *vg* / *L* which, compared to wild type, increased about four-fold in the bik1 mutant and decreased about two-fold in the kip3 mutant (Figure 4B). The catastrophe rate for *S. cerevisiae* and *A. gossypii* cMTs are similar but the growth and shrinkage speeds are three-fold slower in *S. cerevisiae* (Figure 4B) according to averaged published data (Tirnauer *et al.*, 1999; Adames and Cooper, 2000; Kosco *et al.*, 2001; Huang and Huffaker, 2006; Wolyniak *et al.*, 2006; Caudron *et al.*, 2008).

Simulations with 3 cMTs and 6 cMTs were computed, and videos with 3 cMTs are documented for bik1∆ (Video 5), kip3∆ (Video 6) and *S. cerevisiae* cMT dynamic (Video 7, discussed below). Already, the visual representation of one simulation each (Figure 4C) reveals the successful verification of the nuclear mobility in hyphae lacking Bik1 (mainly migration with the simulated flow) or Kip3 (extensive bidirectional movements and bypassing). For quantitative comparisons, number averages for all nuclear movement parameters are compiled in Supplemental Table S2. Box-plot diagrams (Figure 4D) representing 200 simulations each for the mutants and for wild type confirm in a quantitative manner the decrease in active nuclear movements in the absence of Bik1 (shorter cMTs) and the increase in active nuclear movements in the absence of Kip3 (longer cMTs). Similar box-plot diagrams for simulations with 6 cMTs, shown in Supplemental Figure S2A, confirm these conclusions. Interestingly, nuclear movements increased monotonously as a function of the average cMT length (bik1∆ < “wild-type” < kip3∆). Between the two extremes (rare contact of nuclear cMTs with cortical Num1 in the bik1 mutant and more contact than wild type in the kip3 mutant), the average duration of forward and backward movements increased roughly 1.5 and 3.5-fold, respectively (Figure 4D). The frequency of bypassing events also increased (6.4 fold with 3 cMTs and 8.7-fold with 6 cMTs) and the frequency of tumbling events dropped about 2-fold (Figure 4D; Supplemental Figure S2A). The flow speed of 0.009 μm/s used in the wild type simulations (Figure 3) was also applied in these mutant simulations. As the average cytoplasmic flow speed of the *in vivo* experiments was 0.013 μm/s (Grava and Philippsen, 2010), we repeated all simulations implementing this flow speed. The results were similar except for an even higher reduction in backward movements in the simulated bik1 mutant, most likely due to the increased flow speed (see Supplemental Table S2). Altogether, these calculations demonstrate that the sole modification of cMT dynamics is sufficient to explain the phenotypes observed in bik1∆ and kip3∆ mutants, illustrating that cMT dynamics is a key parameter governing nuclear motions in hyphae.

The visualization of the simulated nuclear movements controlled by *S. cerevisiae* cMT dynamic (Figure 4C, right panel) shows a rather passive moving pattern of the five nuclei which seem to mainly migrate with the applied flow speed. We computed 200 simulations (1000 nuclei) with 3 cMTs nucleated per nucleus (Video 7), as well as another set with 6 cMTs per nucleus, and quantified the movements (see summary in Supplemental Table S2). Nuclei dynamic parameters are very similar to those observed in the simulated bik1 Δ mutant, in which cMTs are also short and rarely reach a cortical activator to induce active movements. However, the *S. cerevisiae* simulation reveals longer lasting backward movements and a still low but higher frequency of bypassing events compared to the bik1 mutant. We had performed this experiment knowing that *A. gossypii* evolved from a *S. cerevisiae-like* ancestor, and we thus had wondered to which extend the adaptation of cMT dynamics has been important in switching from a budding mechanism to filamentous growth. The simulation indicates that an increase of the cMT growth rate, possibly through the adjusted influence of MT-associated proteins, without changing the catastrophe rate could have been sufficient to induce movements of nuclei against the cytoplasmic stream and also nuclear bypassing. This scenario, however, would only have worked with a simultaneous ten-fold density increase of cortical Num1 patches as demonstrated in the next chapter.

### Nuclear movements require an adapted density of cortical anchors

The minus-end directed MT motor dynein is responsible for all active nuclear movements in *A. gossypii.* It localizes to −ends and +ends of cMTs and also along cMTs. Hypha with inactivated dynactin or Num1 show severely reduced active nuclear movements concomitant with enrichment of dynein at cMT +ends (Grava *et al.*, 2011). These phenotypes and the observed onset of pulling, when a cMT slides along the hyphal cortex, described by the same authors, support a mechanism for cortical pulling known from studies in *S. cerevisiae.* The dynein/dynactin complex is thought to be inactive while transported either by the kinesin Kip2 towards the cMT +ends or directly recruited to +ends from the cytoplasm (Markus et al., Current Biology 2009; Markus and Lee, 2011; Roberts et al., 2014). When the cMT +end associates with a cortical Num1 patch during sliding at the plasma membrane, the dynein/dynactin complex would be off-loaded, anchored with the amino terminal tail of Dyn1, the dynein heavy chain, to the amino terminal binding domain of Num1, and the dynein motor activated to eventually pull on the cMT on which it was previously hitchhiking (Lee *et al.*, 2003; Moore *et al.*, 2009; Markus and Lee, 2011). Recently, it was demonstrated that dynein is directly switched on by the cortical anchor Num1 (Lammers and Markus, 2015).

The dynein-binding domain and the lipid-binding domain of ScNum1 are conserved in AgNum1, and recently patches of an AgNum1-GFP fusion were observed at the *A. gossypii* cortex (Doris Landmann, unpublished). Knowing the close evolutionary relation between *S. cerevisiae* and *A. gossypii,* we assumed a similar role of Num1 in anchoring and activating dynein in both organisms. As the cortical density of Num1 patches is presently unknown, we ran simulations with different numbers of randomly distributed cortical anchors. As already described above, we envisage that these anchors will engage cMTs that are within their reach, and immediately recruit dynein. The duration of a pulling event is then determined by the off-rate of the motor domain to the cMT (see Materials and Methods). In terms of nuclear movements, this simplified model is sufficient because forces are only created when dynein is anchored at the cortex and engaged with a cMT. With this model, the density of anchors on the cortex (~0.7 anchors per μm^2^ in all simulations discussed so far) is the key parameter controlling the efficacy of dynein-mediated cortical pulling. Hence varying the absolute number of anchors in the simulation can be qualitatively compared to the phenotypes of changing the overall dynein expression level in live cells.

Experimentally reducing the expression level of the dynein heavy chain by truncating the *AgDYN1* promoter decreases oscillatory and bypassing movement frequencies (Grava *et al.*, 2011). In simulations, reducing the amounts of anchors 2-fold (Video 8) or 10-fold (Video 9) clearly reduced active nuclear movements (see also Figure 5A and Supplemental Table S3). The reduction in density also increases the movement frequency ratio (Figure 5B, top; by favoring forward movements induced by the cytoplasmic flow), decreases the frequency of bypassing events (Figure 5B, middle) and increases the frequency of tumbling events (Figure 5B, bottom), as previously described *in vivo* for the prom 180-DYN1 and prom130-DYN1 strains, in which the *DYN1* promoter was shortened to 180 or 130-bp of the original sequence upstream of the start codon thereby reducing gene expression (Grava *et al.*, 2011). We then continuously varied the anchor density from 0 to 1000 molecules per 30 μm of hypha (0 to 3.5 anchors per μm^2^). Above a certain threshold of anchor density, the pattern of nuclear movements is constant, and determined purely by cMT encounters with cortex-bound Num1 patches. The value of the threshold is lower with 6 cMTs per nucleus (Figure 5C, bottom) than with 3 cMTs per nucleus (Figure 5C, top), indicating that increasing the density of anchors can compensate for a reduced count of cMTs. This can be understood since dynein pulling on a single cMT is sufficient to move a nucleus. In the saturated regime where anchors are in excess, the frequency ratio is close to 1 (Figure 5D, left). However, the value measured experimentally is ~1.6, suggesting that active force generation is indeed limiting *in vivo*. In other words, not every cMT contacting the cortex *in vivo* is pulled by dynein. The same conclusion can be derived from the frequencies of bypassing events: the measured *in vivo* value of ~0.026 /min is only half of the predicted plateau value of 0.05 /min (Figure 5D, right). Altogether, these results show that, despite necessary simplifications, our model has captured how nuclear movements depend on dynein, as measured *in vivo.*

Interestingly, an amount of 20 anchors per hypha segment, which corresponds to 0.07 anchors per μm^2^, also matches the observed density of Num1 patches in *S. cerevisiae* (~8 patches per cell). The fact that at this density only very few movements are produced in our simulation (Figure 5A-B) demonstrates that *A. gossypii* had to adjust the density of Num1 patches to be compatible with hyphal growth. Specifically, our simulations suggest that a 10-fold increase in anchor density would allow enough cMT capture to promote sufficient nuclear movements.

### The impact of cytoplasmic streaming on nuclear movements depends on cMT number

To further explore the role of cMT number on the balance between forward and backward movements, we searched for its importance in the context of varying flow speeds. Indeed, hypha growth speed – and the resulting cytoplasmic flow – was previously shown to impact the balance between forward and backward movements (Lang *et al.*, 2010b). However, the role of cMT number in this balance has not been investigated yet. We therefore ran simulations with flow speeds randomly varying between 0 and 0.02 μm/s, corresponding to the range of hypha growth speeds for which *in vivo* data were previously collected (Lang *et al.*, 2010b). The cMT length was controlled as in wild type. For either condition, with 3 or 6 cMTs per nucleus, we generated 200 random values for the flow speed and ran 3 simulations for each of them. For both conditions, the frequency of forward movements was roughly independent of flow speed (frequencies being lower with 6 cMTs, on average) (Figure 6A) whereas backward movements decreased with an increasing flow speed, and this decrease was more pronounced if the nuclei nucleated 3 cMTs (Figure 6B). The movement frequencies ratio thus remains quite stable for nuclei nucleating 6 cMTs but gets higher and highly biased towards forward movements with increasing flow speed for nuclei nucleating 3 cMTs (Figure 6C). Noticeably, the values of frequencies and ratio observed in our simulations are in agreement with the values measured experimentally, once these are plotted according to the naturally occurring variable hypha growth rates (Figure 6A-C, blue dots). Visualizations of nuclear movement patterns for both conditions at a flow speed of 0 and 0.02 μm/s are presented in Figure 6D and 6E, respectively. These patterns show that at a flow speed of 0 μm/s there is no bias for any direction of movement (Figure 6D; 3 cMTs: Video 10; 6 cMTs: Video 11). However, at a flow speed of 0.02 μm/s, a clear bias for forward movements is visible, more importantly for nuclei nucleating 3 cMTs (Figure 6E; 3 cMTs: Video 12; 6 cMTs: Video 13). Is this clear difference between 3 and 6 cMTs per nucleus also seen when flow speeds are tested in hyphae with altered average cMT lengths, like in the bik1 or kip3 mutants? The answer is yes, when cMTs are slightly longer as in the kip3 mutant, and no, when the cMTs are too short in the bik1 mutant, either with 3 cMTs (Supplemental Figure S2B) or 6 cMTs (Supplemental Figure S2C).

Finally, because hyphal growth can reach speeds as high as 0.05 μm/s, we wondered if the reduced bias for increased cMT number would still be true in such extreme cases. We thus ran additional simulations with flow speeds now randomly varying between 0 and 0.05 μm/s. For either condition, with 3 or 6 cMTs per nucleus, we generated 500 random values for the flow speed and ran 3 simulations for each of them. Interestingly, the movement frequencies ratios still remained much lower for nuclei nucleating 6 cMTs than for nuclei nucleating 3 cMTs even at very high flow speeds (Figure 6F). Altogether, these results suggest that having additional cMTs may reduce the bias for forward movements that occurs at rapid hyphal growth, i.e. large flow speeds. This also argues for an evolutionary advantage of spending more time in the G2 phase of the nuclear cycle, as compared to a *S. cerevisiae-like* ancestor, allowing the nuclei to carry duplicated side-by-side SPBs for a longer time than in *S. cerevisiae* (Jaspersen and Winey, 2004).

### Organelle crowding interferes with nuclear movements

The cytoplasm of *A. gossypii* hyphae is a crowded environment (Figure 3D), and in particular larger organelles may hinder the movements of the nuclei. We ran three sets of simulations to test the influence of organelles (Figure 7). In the first set, hyphae were lacking simulated organelles (Figure 7A). In the second set, mitochondria occupying 8.8% of the cytoplasm (Gibeaux *et al.*, 2013) were represented as 550 (orange) spheres of 0.4 μm in diameter, and other large organelles (mainly vacuoles) occupying 2.5% of the cytoplasm (Gibeaux *et al.*, 2013) were represented as 50 (blue) spheres of 0.6 μm in diameter. This condition that is derived from electron tomography measurements is referred to as “wild-type” and was used for all simulations of this publication. In the third set, referred to as “highly crowded”, “mitochondria” were unaltered but the diameter of the “vacuoles” was inflated to 1.4 μm (Figure 7A) simulating the condition in older hyphae (Walther and Wendland, 2004). A soft excluded volume interaction is present between all objects (see Materials and Methods). Similar results were obtained for simulations with either 3 cMTs (Figure 7B-C; Video 14-15) or 6 cMTs (Supplemental Figure S3A-B) per nucleus. The averages for nuclear motility parameters determined from these simulations are summarized in Supplemental Table S4. Between the “empty” and the “highly crowded” simulations, the average duration of forward movements was reduced by ~10% (the ratios were 1.16 with 3 cMTs and 1.10 with 6 cMTs). Crowding also reduced the average duration of backward movements by ~5% (ratios were 1.05 and 1.04), reduced the frequency of bypassing events (ratios 1.05 and 1.30) and increased the frequency of tumbling events by ~20% (ratios 1.18 and 1.17). All these differences are small, which may indicate that the load on dynein motors is only a small fraction of their stall force, because the viscous drag on the nucleus and impeding surrounding objects do not provide much resistance. We next asked if, as for cMT number, the presence of organelles could also change the balance of movements as a function of hypha growth speed. We thus ran simulations with random flow speeds as described for Figure 6, with no or high crowding. For the two conditions, the curves representing the frequencies of forward and backward movements and their ratio, overlapped for all values of the cytoplasmic flow speed (Figure 7C; Supplemental Figure S3B). Altogether, these computations show that the larger organelles affect nuclear dynamics. Unfortunately, the very likely additional influence of the different forms of the endoplasmic reticulum (ER) could not be investigated. ER structures close to the nuclear envelop, within hyphae and close to the cortex were observed by electron tomography (Gibeaux *et al.*, 2013) but the ER network is complex as shown in budding yeast (West *et al.*, 2011) and could not be modelled simply.

**FIGURE 7:**
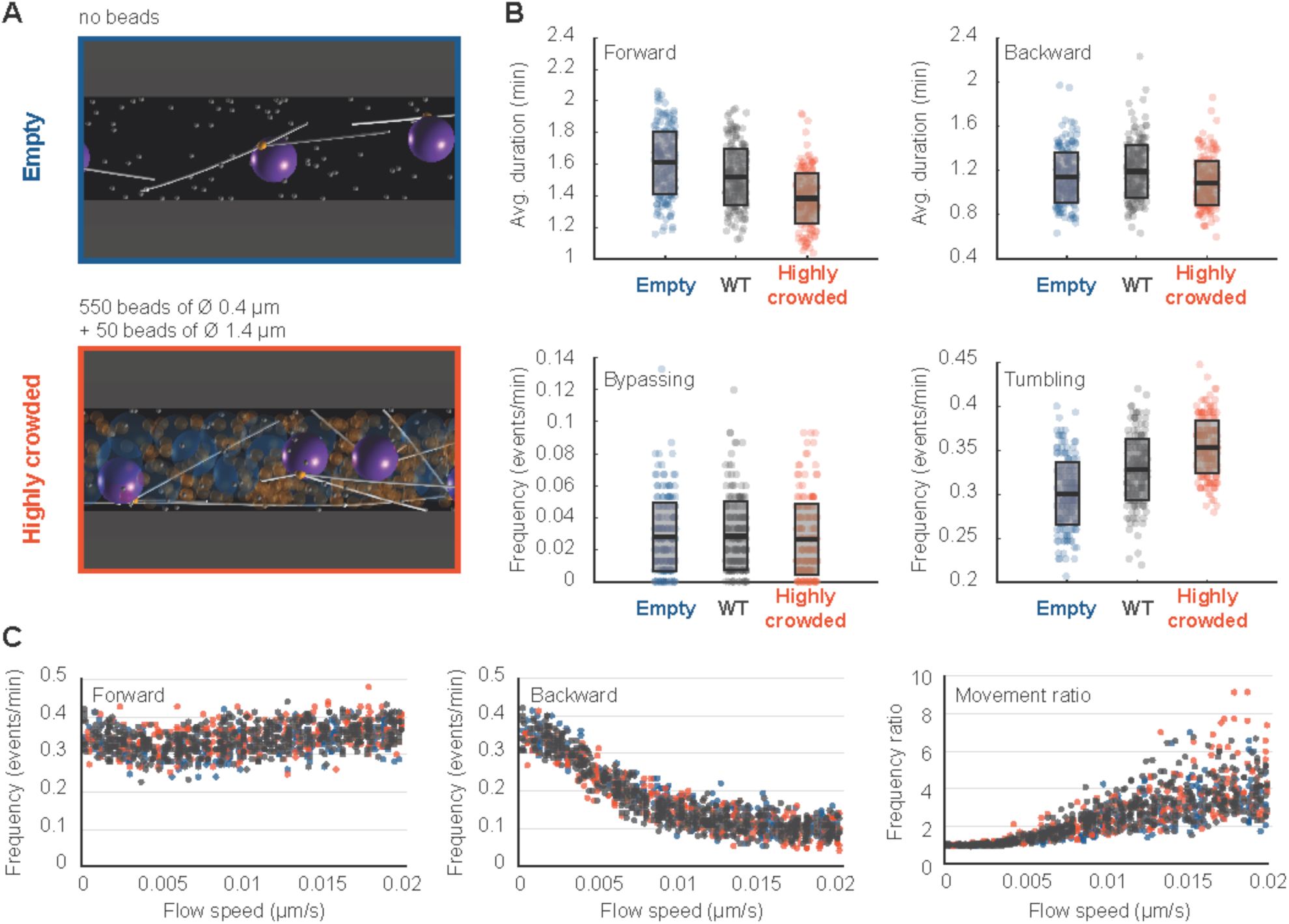
Influence of organelle crowding on nuclear dynamics. **(A)** Snapshots of simulations with uncrowded and highly crowded cytoplasm. To represent the “wild-type” (‘WT’) situation, the simulation included 550 “mitochondria” spheres of 0.4 μm in diameter and 50 “vacuoles” spheres of diameter 0.6 μm. These objects are removed in the un-crowded ‘empty’ case (top), while in the highly crowded case (bottom), the 50 “vacuoles” were enlarged to a diameter of 1.4 μm. **(B)** Boxplots of nuclear dynamics measurements comparing the simulated ‘WT’ situation to ‘empty’ and ‘highly crowded’ simulations. The average duration of forward movements (top left), the average duration of backward movements (top right), the frequency of bypassing events (bottom left) and the frequency of tumbling events (bottom right) are plotted. The ‘empty’, ‘WT’ and ‘highly crowded’ data are plotted in blue, gray and orange, respectively. Circles stand for individual simulations; the thick black line marks the average value and the transparent gray box the standard deviation. **(C)** Plots of the frequencies of forward (left) and backward (middle) movements and their ratio (right) as a function of increasing cytoplasmic flow (from 0 to 0.02 μm/s). Colors as in panel B.

## DISCUSSION

Prior genetic and live-imaging studies had shown that cMT factors and the dynein motor protein were necessary for the active movements of nuclei in multinucleated hyphae of *A. gossypii.* From these data, it was possible to hypothesize that the oscillatory movements of nuclei in growing hyphae of *A. gossypii* had evolved from the *S. cerevisiae* Dyn 1 -Num 1 pathway essential for the pulling of the anaphase spindle into the bud concomitant with its forth and back pulling through the bud neck. This now seems to be a viable hypothesis. By simulating the process from first principles, we here demonstrated that the pulling action of cortically anchored dynein motors on cMTs originating from the SPB is sufficient to explain active nuclear movements observed *in vivo* in a quantitative manner. Importantly, the obtained agreement between experiments and models was not achieved by adjusting various parameters to fit the desired behavior. Instead, the key parameters of the model, except one, had been previously determined experimentally (see Table 2), including the diameter of the cell, the densities and dimensions of the nuclei and other objects (Gibeaux *et al.*, 2013), the number of cMTs per nuclei (Gibeaux *et al.*, 2012), the growth rate at the plus-ends, and MT lengths in the mutants (Grava and Philippsen, 2010). These latest values allowed us to estimate the catastrophe rate by fitting the length distribution. We could use the motile and force parameters of yeast dynein, which had been measured in single molecules biophysical studies. The only unknown biological parameter was the density of cortical anchors in the simulation, which encapsulates unknow factors such as the concentration of dynein molecules in the cell, and the effectiveness of the transport, activation and anchoring mechanisms. We therefore explored the effect of this parameter systematically (Figure 5). As expected, reducing the quantity of anchors in the cell directly reduced or disabled nuclear movements, but increasing anchor density quickly led to a plateau where every cMT contacting the cortex found at least one force generator (Figure 5C). We selected an intermediate value, which allowed us to fit all the experimental data (see Figure 3 and summary of the quantifications in Table 1). The model reproduced the leading position of the SPB on the nucleus during the movements (see Video 4) and recapitulated the rates of forward and backward movements observed for different values of the cytoplasmic flow, both qualitatively and quantitatively (Figure 6). By comparing the observed motion of the nuclei, and the dependence in the simulation as a function of the density of cortical anchors, we concluded that not every cMT contacting the cortex would be pulled by dynein (Figure 5). This indicates that cytoplasmic flow and dynein-generated forces both contribute significantly to nuclear motions. This is interesting, because the associated motions are physically of a different nature. Cytoplasmic flow is a convective motion and the total distance travelled is proportional to time *x*~*t 𝑣_flow_*. The dynein-mediated movements however are stochastically directed towards or away from the hyphae tip, producing without flow and at long time scales a diffusive motion characterized by the relation 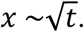 Because dynein easily overcomes the hydrodynamic drag of the nuclei, the motion is purely diffusive also in the presence of flow when cortical anchors are in excess (Figure 5, when the ratio of forward / backward movement frequencies is ~1). It seems more advantageous, biologically, to remain away from this regime, to benefit both from a convective motion that keeps the nuclei equidistant to the growing tip of the hyphae, and from the dynein-mediated active diffusion, that permutes nuclei. This means that the cell must avoid excessive cortical pulling on its cMTs, by keeping the density of anchors below a certain threshold, or via some other mechanism.

The model made some interesting findings that help us reinterpret recent experimental results. Firstly, the simulations matched the *in vivo* observations better with 6 cMTs nucleated per nucleus than with 3 (Table 1). We interpret this to reflect that a high proportion of nuclei carries duplicated side-by-side SPBs, which is consistent with the fact that, although SPBs nucleate 3 cMTs on average (Gibeaux *et al.*, 2012), up to 6 cMTs were observed in live imaging of *GFP-AgTub1* “wild type” cells (Lang *et al.*, 2010b). Secondly, our analysis regarding cMT dynamics (Figure 4) demonstrated that the mutant behavior in bik1∆ and kip3∆ cells could be explained solely by a change in the cMT properties at the plus end (polymerization and catastrophe rate). Importantly, this model also highlighted several adjustments that *A. gossypii* had to undergo to adapt its nuclear movements to hyphal growth from a *S. cerevisiae-like* ancestor. Specifically, cMTs became 4-fold longer, mostly through an increased growth rate, making them able to reach the cortex (Figure 4). Additionally, the density of anchors at the cortex had also to increase, such as by 10-fold as suggested by our simulations, to allow enough cMT capture (Figure 5). Interestingly these adaptations seem enough to provide robustness with respect to the high, and changing, organelle density required for hyphal growth (Figure 7).

By keeping our model minimal, we could thus demonstrate that a small set of ingredients are enough to explain the basis of the nuclear movements observed in the growing hyphae of *A. gossypii.* Nevertheless, some open questions remain that were not addressed. For example, we did not include nuclear division, nor the true elongation of the hyphae, or their branching geometry. Also, the growth of cMTs occurs at a constant speed in our model, but it could be regulated for example by cortical ER, which is substantial in budding yeast (West *et al.*, 2011). The model omitted the formation of septa, because this only occurs in older regions of the hyphae. These processes will all be exciting to simulate in the future, but additional work is required to extend the model and exploit it.

Moreover, a process of nuclear repulsion was recently proposed (Anderson *et al.*, 2013), which leads to the creation of cytoplasmic territories and enables division autonomy in *A. gossypii.* These territories increase in size as a nucleus approaches mitosis. They might be mediated by cMT, as it was suggested earlier that cMTs from neighboring nuclei could interact and repulse each other (Philippsen *et al.*, 2005). However, live cell imaging and high resolution analysis of the cMT cytoskeleton by electron tomography did not reveal such interactions (Lang *et al.*, 2010b; Gibeaux *et al.*, 2012). The mechanism of the repulsion is thus still unknown precluding its implementation in the simulation. Nuclei repulsion in *A. gossypii* is inferred from the shape of the distribution of distances between pairs of adjacent nuclei (Anderson *et al.*, 2013). We thus derived such distributions from the 12 reference live cell movies and 12 simulations (Supplemental Figure S1D). We could confirm that nuclei are not distributed randomly in live hyphae, but are instead separated by a characteristic distance. For the simulations, we found however a distribution that is close to what would be expected if the nuclei were randomly positioned, with a minor indication for nuclear repulsion (Supplemental Figure S1D, see legend for more details).

Nuclear movements were different between 3 or 6 cMTs emanating per nucleus but still remained similar in many aspects, especially in regards to their bypassing frequencies (Table 1). Yet the reported bypassing frequencies are six times higher for a nucleus in G1 than in G2 (Gibeaux *et al.*, 2012). According to our model, the duplication state of the SPB is however not expected to account for this observation. This thus raises a more general and fascinating question: how can a nucleus control its movements while progressing through the nuclear cycle_ Regulating cMT dynamics at the plus-end can lead to direct changes in nuclear behavior (Figure 4; Supplemental Figure S2), but how could different SPBs contained in a common cytoplasm provide variable cMT dynamics_ The challenge is that both the site of force production, and the plus end of the cMT that is pulling a nucleus may be distant from this nucleus. They may be actually located closer to another nucleus, such that any diffusible substance emitted by a nucleus would not target the productive cMT. Nature has found a solution, however: during mitosis in *S. cerevisiae,* for example, proteins are able to localize asymmetrically on the two SPBs, although the cytoplasm is shared. This is the case for Kar9 (Liakopoulos *et al.*, 2003) and Bub2 (Pereira *et al.*, 2000), for instance. Interestingly, the asymmetric localization of Kar9 to one SPB (and MT plus ends) requires a fine regulation through the cyclin-dependent kinase Cdc28. Bik1 binds directly to Kar9 and promotes its phosphorylation, which affects its asymmetric localization to one SPB and associated cMTs (Moore *et al.*, 2006; Moore and Miller, 2007). Moreover, it has also been shown that the XMAP215 homologue Stu2, localizes on the SPB and thus regulates the dynamics of the cMTs anchored to it (Usui *et al.*, 2003). It is therefore possible to imagine that a nucleus, by adjusting the state of the SPBs, could control plus-end cMT dynamics throughout the nuclear cycle and therefore its movements. However, whether this is the case in the multinucleated hyphae of *A. gossypii,* has not been investigated yet. It is actually relevant to note, that dynein is symmetrically distributed to pre-anaphase SPBs of *A. gossypii* (Grava *et al,* 2011; Grava and Philippsen, unpublished) whereas asymmetric distribution has been reported for the pre-anaphase SPBs in *S. cerevisiae* (Grava *et al.*, 2006). Hence, investigating the nature of mechanisms able to regulate the motile machinery as a function of the nucleus-cycle will be an exciting task for future research. Finally, our study highlights that, beyond its usefulness to validate a particular model, modular software such as *Cytosim* can be used to simulate the cytoskeleton in many different configurations, and thus offers a way to unify our understanding of nuclear migration across the Eukaryotic kingdoms.

## MATERIALS AND METHODS

### Extraction of Nuclei 2D coordinates from live imaging data

We used 3 recordings of live-hyphae expressing GFP-tagged histone H4 (see Supplemental Table S1 and Video 1). The methods used to generate these movies are described in corresponding references. From 12 hyphae recorded over 25-30 min, we extracted the 2D coordinates of the hypha tip and that of the first 5 proximal nuclei (and their progeny). Tracking was done in Fiji (Schindelin *et al.*, 2012) using the tracking plugin from Fabrice Cordelières (Institut Curie, France). Coordinates were exported as a text file and further processed with MATLAB (The MathWorks, Inc.). The vector described by the coordinates of the hypha basis and tip was used to apply a rotational matrix to the data points along the hypha axis so that all hyphae can be uniformly analyzed with each other and with the simulations. Except for the analysis presented in Supplemental Figure S1, we smoothed the trajectories using a Savitzky-Golay filter (smooth function, ‘sgolay’ method, default span and degree) to account for pixel size derived inaccuracy of tracking resulting in artefactual nuclear displacements. The average growth speed of each hypha was calculated from the tip positions at the start and end of the movies, and the resulting value was used as a threshold for the classification of the nuclear movements.

### First principles modeling of nuclear dynamics

The model of nuclear movements in *Ashbya gossypii* was developed using *Cytosim,* a stochastic engine that can simulate flexible cytoskeletal fibers, diffusible particles, and other objects in a confined environment (Nedelec and Foethke, 2007). *Cytosim* is based on Brownian dynamics as well as on a stochastic description of the most relevant microscopic processes, such as binding/unbinding of molecules. The motion of a sphere located at position x is defined by an over-damped Langevin equation: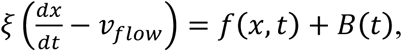 where the right-hand terms are the deterministic and random forces respectively, and *ξ* is a drag coefficient calculated with Stokes law from the viscosity and the size of the sphere. Such equations are suitable because they accurately describe the motion of small objects in a viscous fluid, characterized by a very low Reynolds number (Nedelec and Foethke, 2007). The equations for filaments are similar, and were given previously (Nedelec and Foethke, 2007). A soft excluded volume interaction limits the overlapping of the objects in the simulation. If two spheres of radius ***R*** and ***R’*** are at position ***x*** and ***x’*** such that |*x* + *x* ′| < *R* + *R′* a Hookean force of resting length *R* + *R′* and stiffness 100 pN/μm is applied between the two centers. Similarly, the objects are also confined within the cylinder by forces that are always perpendicular to the cylinder, corresponding to frictionless boundaries. For simplicity, the hypha cell is simulated as a cylinder with periodic boundaries, a standard approach to simulate an unbounded system. Its fluid is in uniform motion and drags along all objects within, but we neglected the variations of flow along the hyphae (Supplemental Figure S1C), since this would not be consistent with periodic boundaries. We also neglected the Poiseuille flow profile that is expected in the transverse section of the hypha and also all kinds of hydrodynamic interactions, by which the motion of a nucleus could affect nearby nuclei. Cortical anchors are randomly positioned on the cylinder surface, and remain immobile. An anchor can engage any cMT that comes within its capture radius (75 nm) at a specified rate (5 s^-1^). Engaged anchors immediately recruit a dynein force generator, which is governed by a linear force-velocity relationship: they move near their maximum speed (25 nm/s) towards the minus-end if the resting force is small, and stall at a force of 7 pN. The duration of the interaction is determined by the exerted force with detachment rate constant 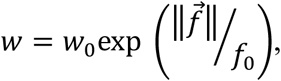 with *f*_0_ = 7 pN and *w*_0_ = 0.64 s^-1^. At the start of the simulations, 5 nuclei are randomly positioned within the hypha, and spheres are added randomly to mimic volume occupancy due to large cytoplasmic organelles. We used 550 beads of 0.4 μm in diameter to fill 8.8% of the cytoplasm and 50 beads of 0.6 μm in diameter for 2.5%, as occupied by mitochondria and other large spherical organelles, respectively (Gibeaux *et al.*, 2013). The nuclei are spherical, and one point with 3 or 6 cMT *nucleators* on their surface represents the SPB. Each nucleator can generate one cMT, and will stay attached and inactive to the minus end of this cMT, which is not dynamic. Cytoplasmic MTs are thus anchored through their minus ends to the SPB, allowing unrestricted rotational freedom so that they can orient in all directions as previously observed (Lang *et al.*, 2010b; Gibeaux *et al.*, 2012). Cytoplasmic MTs grow – and stochastically switch to shrinkage – from their plus ends at a constant speed and with dynamics previously described (Grava and Philippsen, 2010). When a cMT eventually disappears, it is replaced by a new one with a nucleation time of 1s, such that the number of cMT per SPB is nearly always maximal. This new cMT is created with a random orientation. The number of cMT in the simulation is thus either 3 or 6, corresponding to the average number of cMT measured in cells for isolated or paired SPBs. We have not included the variability around the mean in the number of cMT, to better observe the difference between the two conditions. The key physical parameters of the model have been obtained from the literature (see Table 2). Numerical parameters are appropriately chosen to ensure sufficient precision (for example, the time step was 50 ms). The MT assembly dynamics requires ~100s to equilibrate (this time is determined by the catastrophe time), while steric overlaps are resolved much faster. For extra safety, we skipped 600s from the initial configuration, and recorded the trajectories of the nuclei for 1800 seconds (30 min) at 2 s intervals. From these records, the 3D coordinates at 30 s intervals (to match the experimental data) and the flow speed (to be used as a threshold for the movement analysis) were directly processed with MATLAB for analysis. Cytosim is an Open Source project (www.github.com/nedelec/cytosim) and the configuration file to run the wild-type simulation is provided as supplementary material.

### Automated analysis of nuclei dynamics

Nuclei coordinates along time from both experimental and simulation sources were analyzed similarly using MATLAB. Minor differences in the analysis were imposed by the fact that experimental data were effectively 2D, whereas the simulation data were 3D, with periodic boundaries limits. For individual nucleus, speeds and directions were calculated at each time point. From this and the flow speed (obtained from hypha growth speed or set parameter in the simulation and used as threshold), the durations of forward, backward and tumbling events were calculated. The frequency, average duration and average speed (i.e. average of the instantaneous velocities) of these events were then deducted. The x-position offset from the center of one nucleus to any other one were calculated and a change in sign was used to indicate a bypassing event. The frequency of these events was deducted. Statistical tests and plots were performed in Microsoft Excel or MATLAB (Box plots were generated using Rob Campbell’s notBoxPlot function in MATLAB). All MATLAB scripts written for this article can be obtained upon request.

## ACKNOWLEDGMENTS

We thank the members of the Nédélec group for helpful discussions and support, and especially all the people who have been involved, past and present, in the development of *Cytosim.* We acknowledge EMBL services for maintaining IT infrastructure, and particularly the computer cluster. We are grateful to Sandrine Grava for providing raw data of microtubule length measured in *A. gossypii* mutants, to Hans-Peter Schmitz and Doris Nordmann for AgNum1-GFP data, and to Yves Barral for helpful discussions. We also thank Rebecca Heald for her support, Christian Häring and Julio Belmonte for critical reading.

FN, RG and AZP designed the overall project. RG and PP raised the main biological questions. FN and AZP further developed the *Cytosim* platform to implement the model. PP provided the supporting experimental data and his laboratory’s expertise. RG ran all the simulations and did the analysis, supported by FN and AZP. RG, PP and FN wrote the manuscript with inputs from AZP. All authors discussed the results and commented on the manuscript.

## SUPPLEMENTAL FIGURES

**SUPPLEMENTAL FIGURE S1:**
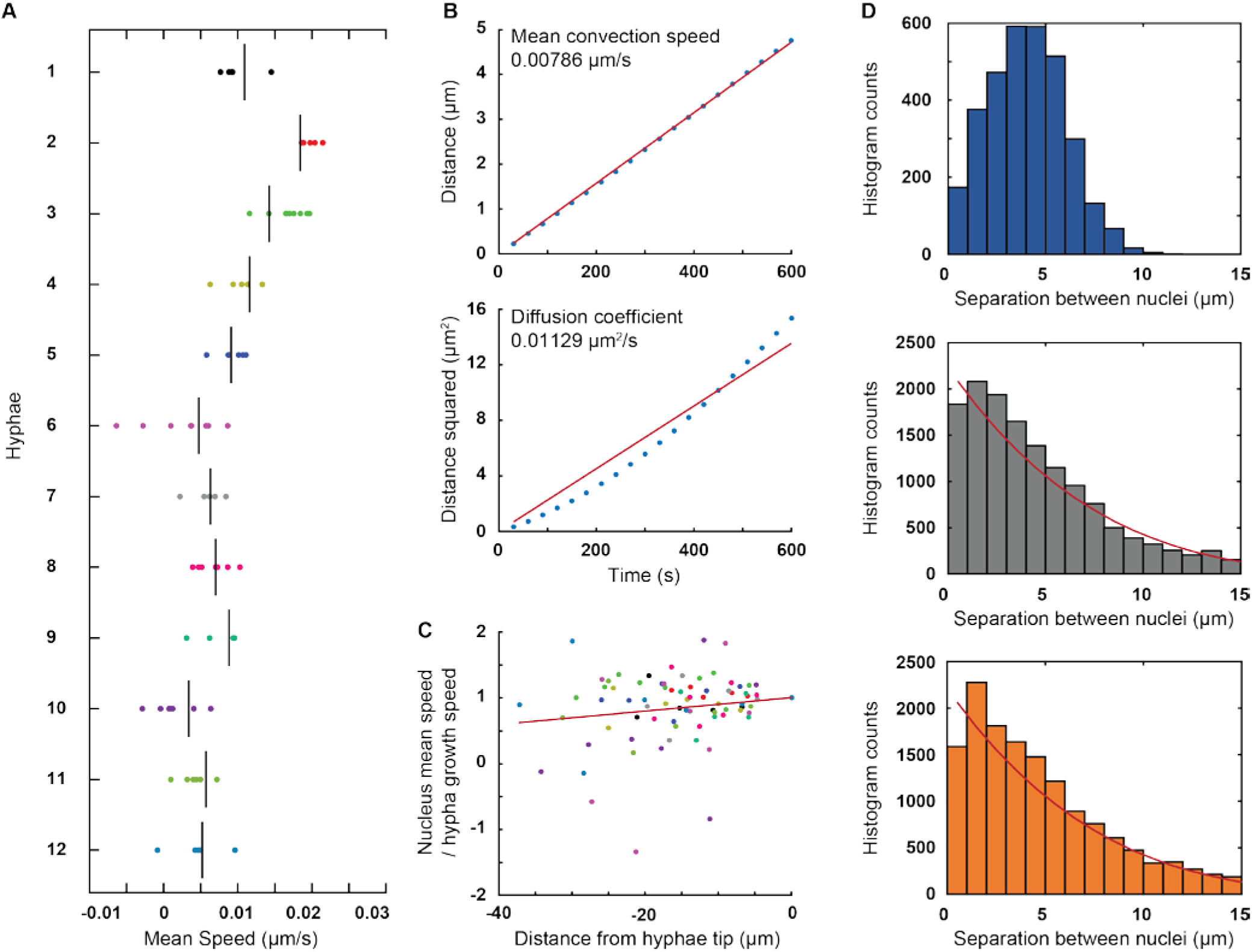
General characteristics of nuclei movements in hyphae. The main axis of the hyphae was calculated by a principal component analysis using the coordinates (x, y) of all nuclei and hyphae tip, derived from the tracking. We then used the coordinate x of the nuclei along this axis to calculate global characteristics of nuclei motions, pooling the data from 12 hyphae. **(A)** The average speed of nuclei movement is directed towards the hyphae tip, and is approximately equal to the speed of hyphae growth. For each hypha, each dot indicates the mean speed of motion of one nucleus, while the vertical line indicates the speed of tip growth. **(B)** The mean and the variance of x(t+τ) – x(t) is plotted as a function of the time interval τ, including all time-points t for which this can be calculated. Points indicate the experimental data, while the lines indicate linear fits constrained to cross the origin. The good agreement show that the nuclei move at an average speed of ~0.008μm/s, with a diffusive component of ~0.011 μm^2^/s. **(C)** The speed of a nucleus depends on its distance to the hyphae tip (negative here), with more distant nuclei moving slower. The line indicates the best linear fit minimizing the sum of the squared residuals that crosses (0,1). These results indicate that the cytoplasmic flow is decreasing away from the hyphae’s tip. This effect was however not included in our simulation, since it would not be consistent with the periodic boundary conditions.(**D**) Distribution of the separation between nuclei in live recordings and in simulations. The difference in the x-coordinates (i.e. along the hypha axis) of the nuclei centers, was used. A ‘zero’ value corresponds to side-by-side nuclei at the same position of the hyphae, but not overlapping. Values for all experimental data were plotted in blue (top), for 6 simulations with 3 cMTs in grey (middle) and for 6 simulations with 6 cMTs in orange (bottom). The separation means were of 4.05 ± 1.96 μm, 4.96 ± 4.20 μm and 4.92 ± 3.92 μm, respectively. The red lines represent the outcome of randomly distributing the 5 nuclei. It has a theoretical mean of 5 μm.

**SUPPLEMENTAL FIGURE S2:**
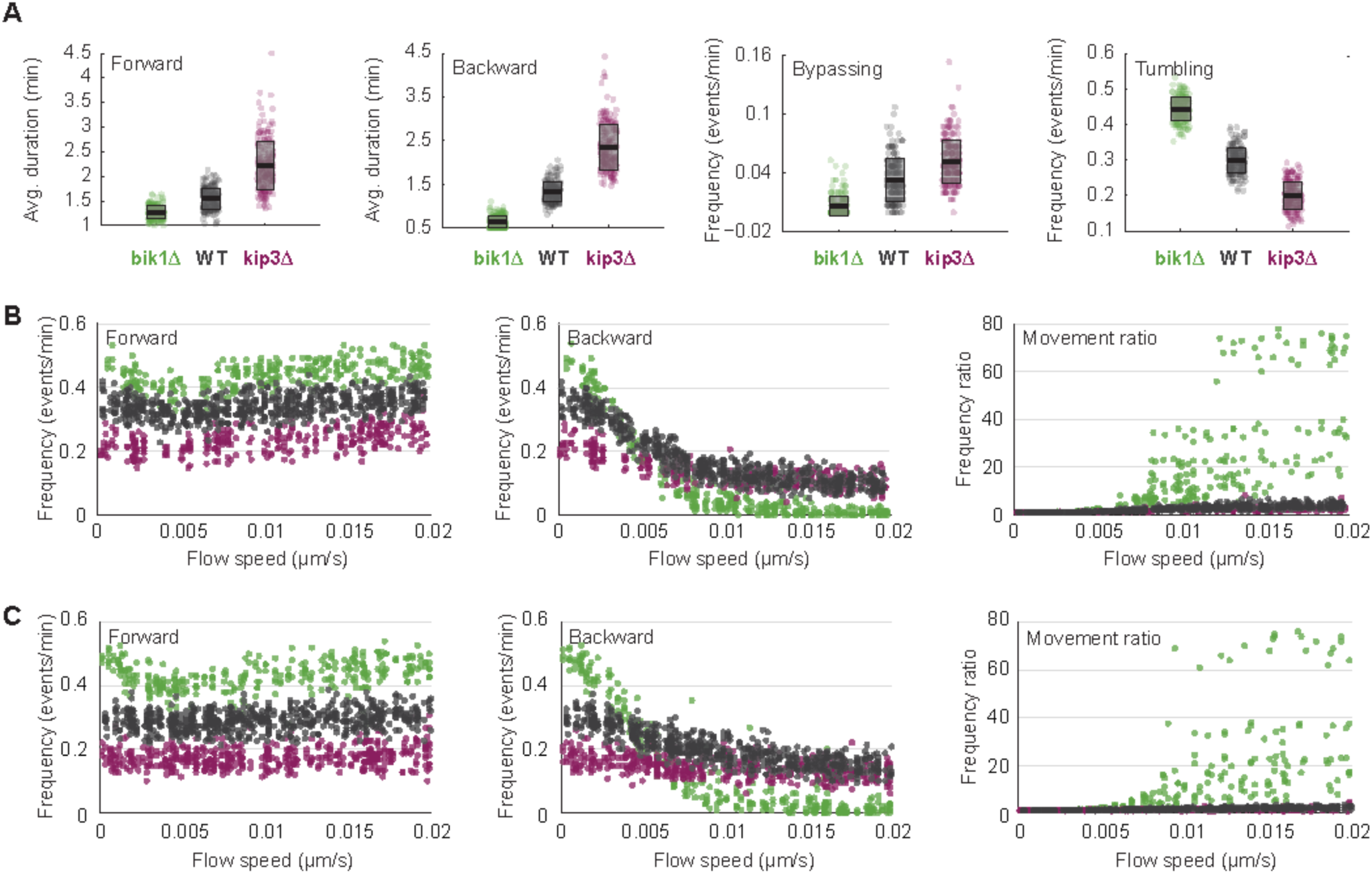
Importance of cMT dynamics in modulating nuclear dynamics. **(A)** Characteristics of nuclear movements in “wild-type” and “mutant” simulations with nuclei nucleating 6 cMTs. The average duration of forward movements (left), the average duration of backward movements (middle left), the frequency of bypassing events (middle right) and the frequency of tumbling events (right) are plotted. Colors as in Figure 4. Circles stand for individual simulations; the thick black line marks the average value and the transparent gray box the standard deviation. **(B)** Plots of the frequencies of forward (left) and backward (middle) movements and their ratio (right) as a function of cytoplasmic flow in “wild-type” and “mutant” simulations with nuclei nucleating 3 cMTs. Colors as in panel A. **(C)** Plots of the frequencies of forward (left) and backward (middle) movements and their ratio (right) as a function of cytoplasmic flow in “wild-type” and “mutant” simulations with nuclei nucleating 6 cMTs. Colors as in panel A.

**SUPPLEMENTAL FIGURE S3:**
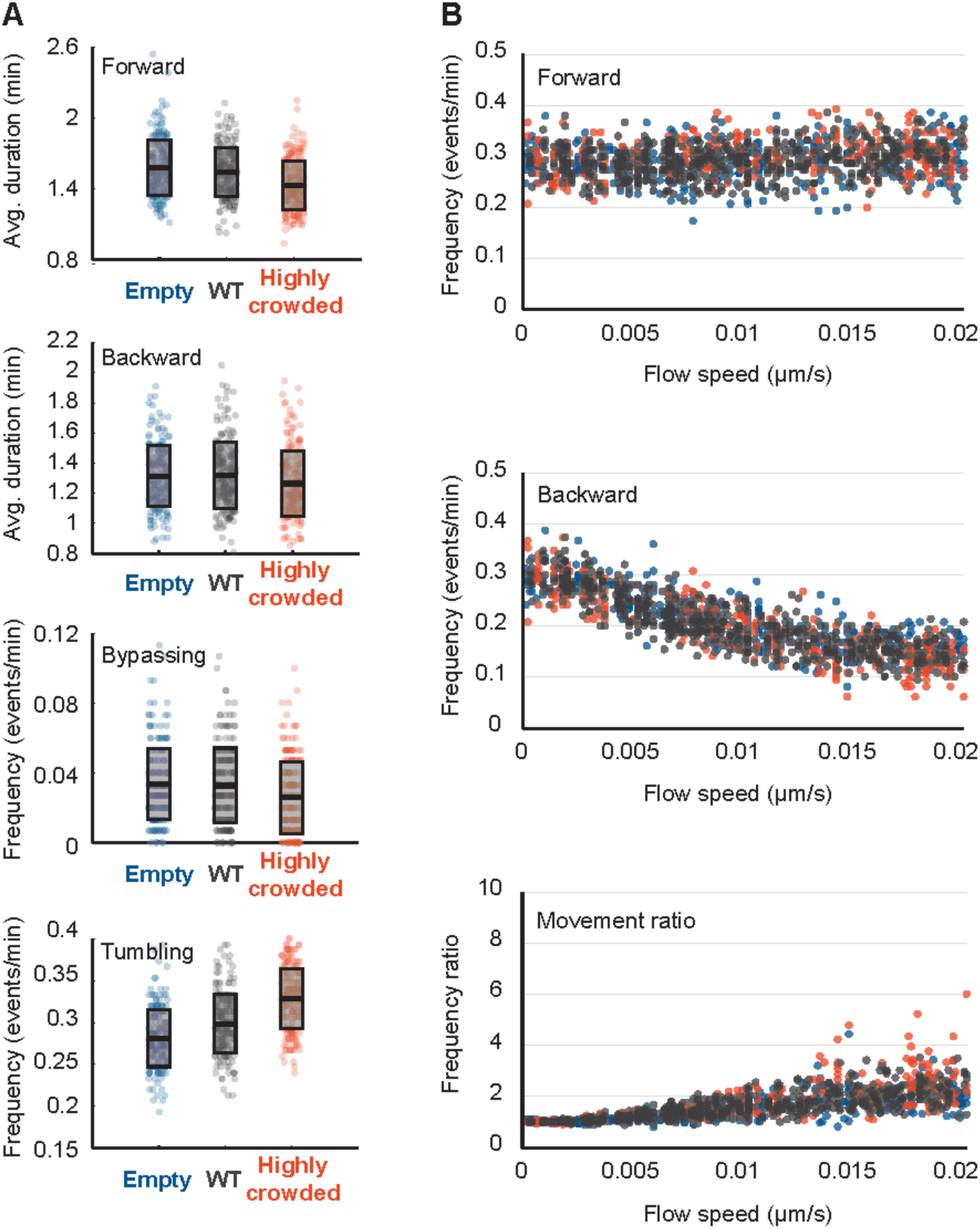
Consequence of organelle crowding on nuclear dynamics in simulations with nuclei nucleating 6 cMTs. **(A)** Characteristics of nuclear movements in the simulated ‘WT’, ‘empty’ and ‘highly crowded’ simulations. The average duration of forward movements (top), the average duration of backward movements (middle top), the frequency of bypassing events (middle bottom) and the frequency of tumbling events (bottom) are plotted. The ‘empty’, ‘WT’ and ‘highly crowded’ data are plotted in blue, gray and orange, respectively. Circles stand for individual simulations; the thick black line marks the average value and the transparent gray box the standard deviation. (**B**) The frequencies of forward (top) and backward (middle) movements and their ratio (bottom) as a function of increasing cytoplasmic flow (from 0 to 0.02 μm/s). Colors as in panel A.

SUPPLEMENTAL TABLES

**Supplementary Table S1.**
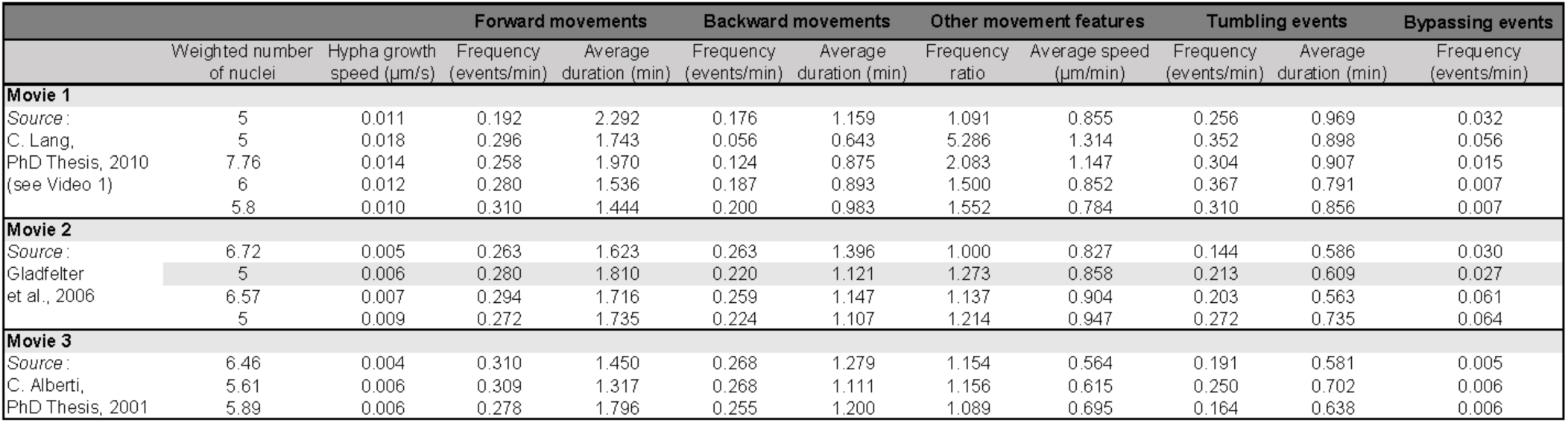
Quantitative description of nuclear dynamics in individual hyphae from live-imaging experimental data.

**Supplementary Table S2.**
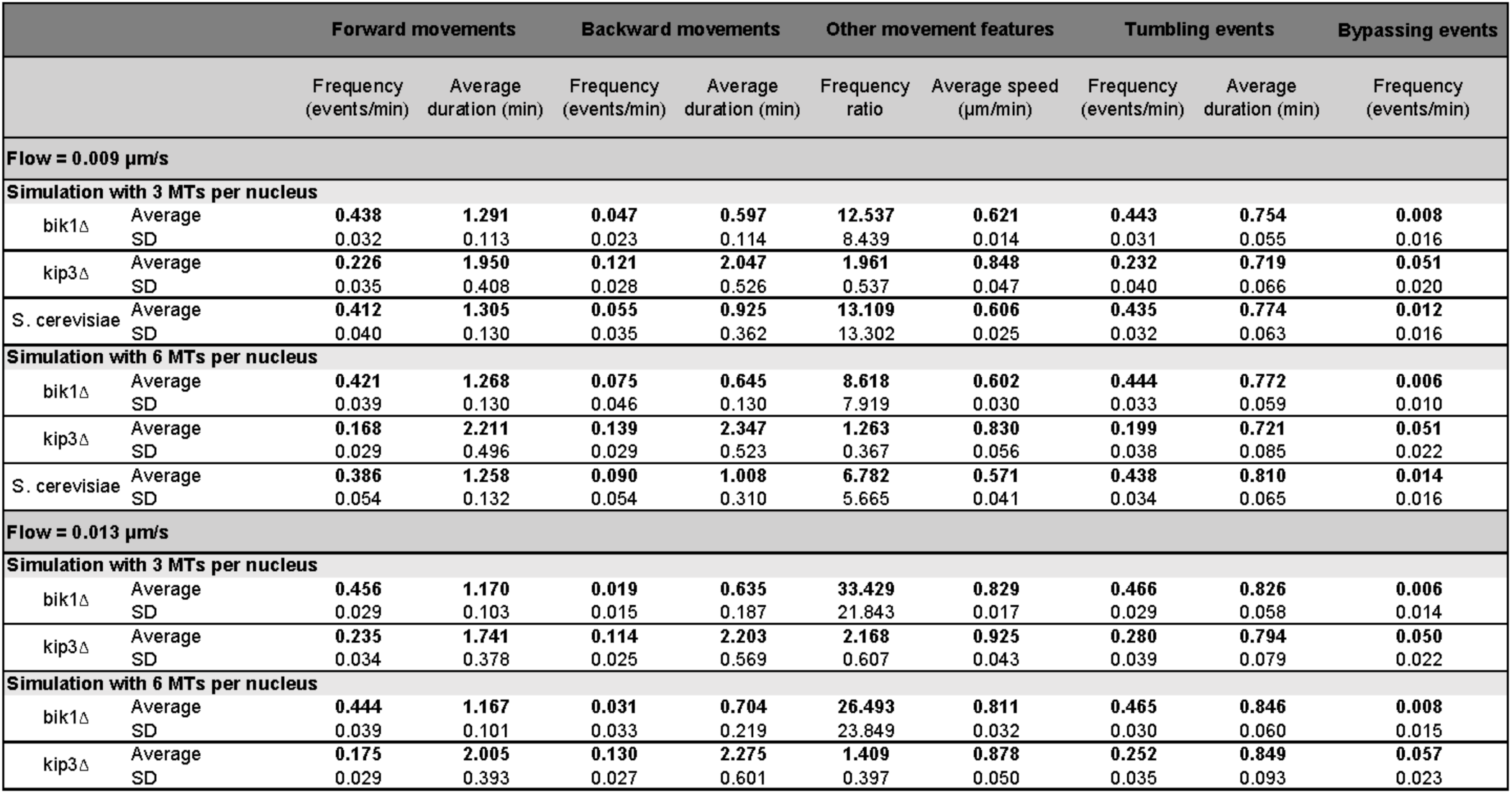
Nuclear dynamics quantification averages from microtubule dynamics simulations.

**SupplementaryTable S3.**
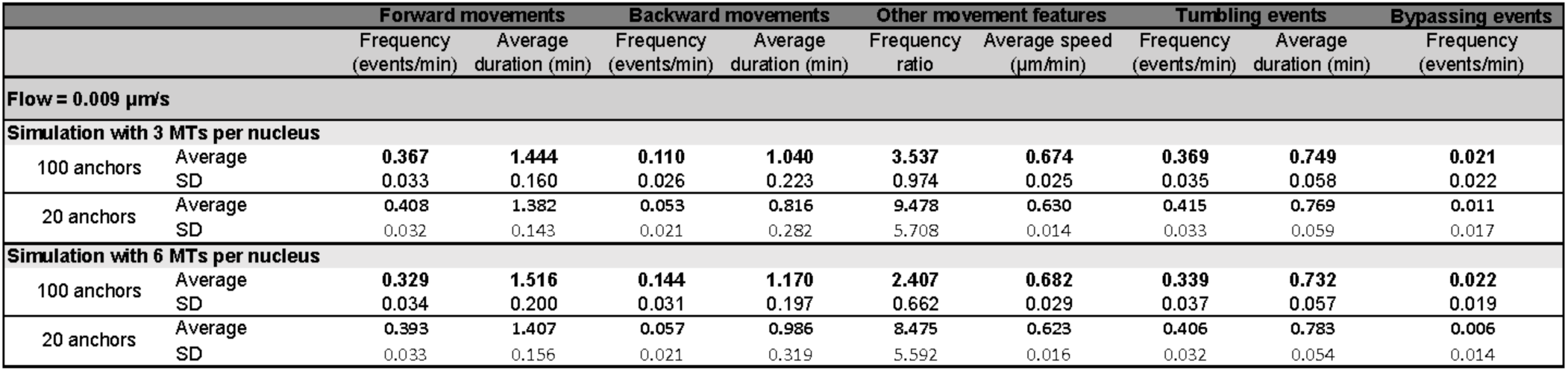
Nuclear dynamics quantification averages with cortical anchors amount decreased to 100 or 20 anchors per hypha

**Supplementary Table S4.**
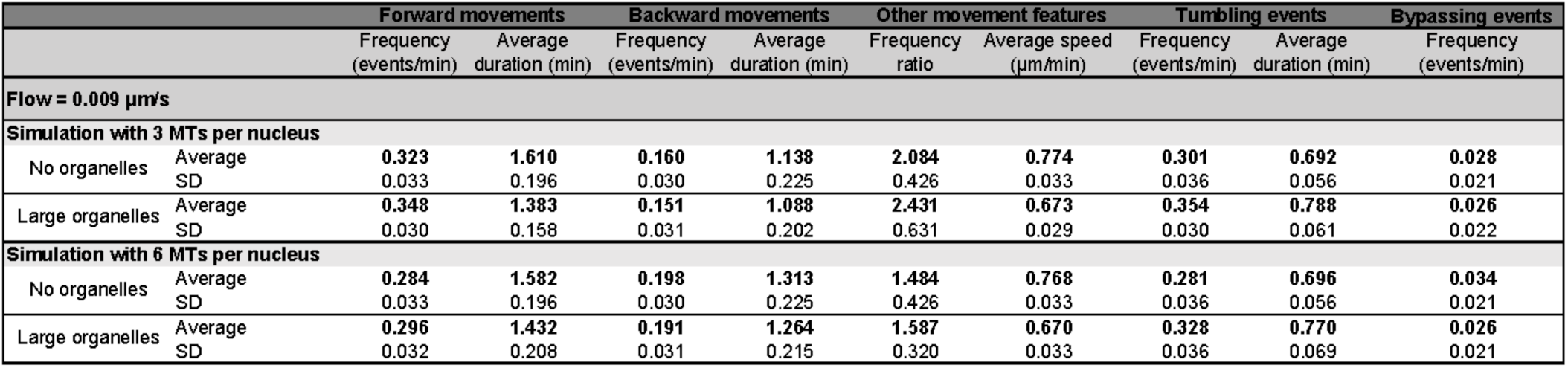
Nuclear dynamics quantification averages with orwithout cytoplasmic organelle crowding.

|  |  | Forward movements |  | Backward movements |  | Other movement features |  | Tumbling events |  | Bypassing events |
| --- | --- | --- | --- | --- | --- | --- | --- | --- | --- | --- |
| | | Frequency<br>(events/min) | Average<br>duration (min) | Frequency<br>(events/min) | Average<br>duration (min) | Frequency<br>ratio | Average speed<br>($\mu\text{m}/\text{min}$) | Frequency<br>(events/min) | Average<br>duration (min) | Frequency<br>(events/min) |
| Flow = 0.009 $\mu\text{m}/\text{s}$ | | | | | | | | | | |
| Simulation with 3 MTs per nucleus |  |  |  |  |  |  |  |  |  |  |
| No organelles | Average | 0.323 | 1.610 | 0.160 | 1.138 | 2.084 | 0.774 | 0.301 | 0.692 | 0.028 |
|  | SD | 0.033 | 0.196 | 0.030 | 0.225 | 0.426 | 0.033 | 0.036 | 0.056 | 0.021 |
| Large organelles | Average | 0.348 | 1.383 | 0.151 | 1.088 | 2.431 | 0.673 | 0.354 | 0.788 | 0.026 |
|  | SD | 0.030 | 0.158 | 0.031 | 0.202 | 0.631 | 0.029 | 0.030 | 0.061 | 0.022 |
| Simulation with 6 MTs per nucleus |  |  |  |  |  |  |  |  |  |  |
| No organelles | Average | 0.284 | 1.582 | 0.198 | 1.313 | 1.484 | 0.768 | 0.281 | 0.696 | 0.034 |
|  | SD | 0.033 | 0.196 | 0.030 | 0.225 | 0.426 | 0.033 | 0.036 | 0.056 | 0.021 |
| Large organelles | Average | 0.296 | 1.432 | 0.191 | 1.264 | 1.587 | 0.670 | 0.328 | 0.770 | 0.026 |
|  | SD | 0.032 | 0.208 | 0.031 | 0.215 | 0.320 | 0.033 | 0.036 | 0.069 | 0.021 |

## VIDEOS

**VIDEO 1:** Example of live-hyphae expressing GFP-tagged histone H4. DIC in gray and GFP signal in green.

**VIDEO 2 to 9:** These 14 movies represent various conditions, but the graphical representation is always identical, unless otherwise stated. The hypha (black) contains 5 nuclei (purple). One SPB (orange) on each nucleus may nucleate either 3 or 6 cMTs (white lines) that are elongating or shortening at the (distal) plus-end. MTs may be pulled by dynein motors that load on fixed anchored at the cortex. The anchor points are displayed gray, and then green if a dynein is engaged. Forces produced by dynein motors and transmitted along the MTs can move the nucleus. The simulation uses periodic boundary conditions, and objects leaving on the right side reappear and the left side, and vice-versa. The cytoplasm is also filled with various organelles. Unless otherwise noted, the simulation contains 550 spheres of diameter 0.4μm (mitochondria, orange) and 50 spheres of diameter 0.6μm (blue, large spherical organelles) and a uniform cytoplasmic flow is moving all objects from left to right at a speed of 0.009 μm/s.

**VIDEO 2:** A “wild-type” simulation with 3 cMTs per nucleus.

**VIDEO 3.** A “wild-type” simulation with 6 cMTs per nucleus.

**VIDEO 4.** A “wild-type” simulation with 3 cMTs per nucleus. The same simulation as shown on Video 1 is displayed here, using a different style. The organelles are not displayed, and the nuclei are semi-transparent such that the SPB (orange) is always visible. The cortical anchors are drawn in dark gray when they are unbound/inactive, and green when they are loaded with bound/active dynein. The re-orientation of the SPB upon pulling is clearly visible, for example for the right-most nucleus during the interval 1400–1700s.

**VIDEO 5.** A simulation with bik1∆–like MT dynamics. Everything is like the “wild-type” with 3 cMT/nucleus, except that MT growth and catastrophe rates were altered to represent the MT dynamics in bik1∆ mutants (see Figure 4A-B).

**VIDEO 6.** A simulation with kip3∆–like MT dynamics. Everything is like the “wild-type” with 3 cMT/nucleus, except that MT growth and catastrophe rates were altered to represent the MT dynamics in bik3∆ mutants (see Figure 4A-B).

**VIDEO 7.** A simulation with *S. cerevisiae*–like MT dynamics. Everything is like the “wild-type” with 3 cMT/nucleus, except that MT growth and shrinkage as well as catastrophe rates were altered to represent the MT dynamics measured in *S. cerevisiae* (see Figure 4A-B).

**VIDEO 8.** A simulation with a reduced number of cortical anchors. Everything is like the “wild-type” with 3 cMT/nucleus, except that the number of anchors on the cortex has been reduced to 100 instead of 200

**VIDEO 9.** A simulation with an even more reduced number of cortical anchors. Everything is like the “wild-type” with 3 cMT/nucleus, except that the number of cortical anchors has been reduced to 20 instead of 200.

**VIDEO 10.** A “wild-type” simulation with 3 cMTs per nucleus and no cytoplasmic flow (speed = 0 μm/s.)

**VIDEO 11.** A “wild-type” simulation with 6 cMTs per nucleus and no cytoplasmic flow (speed = 0 μm/s).

**VIDEO 12.** A “wild-type” simulation with 3 cMTs per nucleus at a high cytoplasmic flow (speed = 0.02 μm/s).

**VIDEO 13.** A “wild-type” simulation with 6 cMTs per nucleus at a high cytoplasmic flow (speed = 0.02 μm/s).

**VIDEO 14.** An “empty” simulation without organelles. The simulation does not contain the extra spheres that are used to represent mitochondria and large spherical organelles.

**VIDEO 15.** A simulation with “high” crowding. The simulation contains 550 spheres of 0.4 μm in diameter (orange, simulating mitochondria) and 50 beads of 1.4 μm in diameter (blue, simulating large spherical organelles in old hyphae).

